# Cover crop microbiomes affect legume cash crop growth but not consistently through enriching nitrogen-fixing rhizobia

**DOI:** 10.1101/2025.10.23.684197

**Authors:** Kayla M. Clouse, Elizabeth L. Paillan, Cody L. DePew, Jennifer E. Harris, Amanda P. Jason, Kelsey Mercurio, Andy L. Swartley, Ellen Bingham, Liana T. Burghardt

## Abstract

Harnessing plant-microbe interactions offers a promising path to reduce chemical inputs and enhance crop resilience in agricultural systems. However, microbial inoculants often fail to persist or function consistently across soils, which limits their broad utility. Here, we explore whether legume cover crops can be used to create microbial legacies that improve nodulation and nitrogen fixation in downstream legume cash crops. In a greenhouse experiment, we inoculated four cash crops with rhizosphere and nodule microbiomes derived from different legume cover crops, then used 16S rRNA and *nifH* amplicon sequencing to profile bacterial and diazotroph communities, respectively. Host identity shaped cover crop rhizosphere and nodule microbial communities, and specific taxa within these communities predicted nodulation and growth in some cash crops. Host specificity varied widely across cash crops, with alfalfa narrowly dependent on a specific symbiont and common bean and fava bean forming more permissive, taxonomically diverse nodule communities. Increased nodulation did not consistently improve biomass, and outcomes in some cash crops depended on more than symbiont compatibility alone. In particular, common bean growth was predicted by both rhizobial and non-rhizobial taxa, while soybean nodulation was shaped by its compatible symbiont as well as a mismatched rhizobial taxon associated with other hosts. Together, these results suggest that cover crops can shape cash crop microbiomes and productivity in host-specific ways, requiring precise symbiont matching in selective hosts but offering more flexible, multi-taxon management opportunities in permissive hosts.

## INTRODUCTION

Legumes are valued worldwide not only as nutrient-rich food crops but also for their ability to enhance soil fertility through biological nitrogen fixation. By forming symbiotic relationships with rhizobial bacteria that fix nitrogen, legumes can reduce reliance on synthetic nitrogen fertilizers and support more sustainable cropping systems (Herridge et al., 2008). The effectiveness of nitrogen fixation, however, depends not only on the presence of legumes but also on the compatibility between the host plant and the rhizobial strains present in the soil or rhizosphere (Burghardt & diCenzo, 2023; Walker et al., 2020). Since rhizobial communities associated with one legume species may not be able to form effective symbioses with other legume species, the choice of preceding leguminous crops in cropping systems may influence the success of subsequent legumes. Selecting upstream cover crops that enrich beneficial rhizobial communities for target cash crops may therefore offer a strategy to improve both crop productivity and sustainable nitrogen management in legume-based systems. Here, we explore whether symbiont legacies left by legume cover crops shape rhizobial community assembly and the efficiency of nitrogen-fixing symbioses in subsequent legume cash crops.

Rhizobia occupy multiple, distinct ecological habitats both within and outside of their legume hosts (Burghardt, 2020; Denison & Kiers, 2011). In root nodules, rhizobia fix atmospheric nitrogen into ammonia in exchange for carbon compounds produced by plant photosynthesis. Outside of nodules, rhizobia persist as free-living populations in the soil and rhizosphere, where they form diverse and dynamic communities (Denison & Kiers, 2004; Poole et al., 2018). These microbial communities may interact with a wide range of legume hosts and often include hundreds of strains capable of forming symbioses within a single field (Mutch & Young, 2004). In addition, these strains compete for access to legume nodules, where successful colonization can lead to substantial increases in population size (Denison & Kiers, 2011). Upon nodule senescence, rhizobia are released back into the soil and can persist across seasons, thereby influencing downstream soil and plant microbial community composition and functionality.

In agricultural systems, legumes are widely used as both cover crops and cash crops. Cover crops are planted between cash crop seasons to provide ecosystem services like weed suppression, erosion control, and pest management (Finney et al., 2017; Kaye et al., 2019). In addition, nitrogen in cover crop biomass is mineralized by soil microbes after termination, making it available to subsequent cash crops. While this nitrogen release from biomass is well established (Liebman et al., 2018; Parr et al., 2011), legume cover crops may also have important legacy effects on nitrogen cycling through the rhizobial populations they enrich. The impact of this legacy depends on host-symbiont specificity: legume cover and cash crop species can differ in their specificity for rhizobial symbionts, with some legumes forming nodules only with a particular bacterial species or subspecies, and others associating with multiple rhizobial genera (Checcucci et al., 2017). For example, alfalfa (*Medicago sativa*) forms nodules with only *Sinorhizobium*, whereas common bean (*Phaseolus vulgaris*) can nodulate with a much wider range of rhizobial genera (Peters et al., 1986; Shamseldin & Velázquez, 2020). The taxonomic identity of resident rhizobia, shaped by prior cover crop use, may therefore determine whether subsequent cash crops can effectively form nodules and fix nitrogen.

The association between legumes and rhizobia has been deliberately managed for agricultural benefit for over a century. By improving nitrogen fixation and soil fertility, this association can reduce fertilizer needs; however, the application of nitrogen fertilizer remains high due to spatial and temporal variability of nitrogen fixation in leguminous crops (Galloway et al., 2008; Herridge et al., 2008). Rhizobial inoculants are increasingly applied in agricultural systems to enhance biological nitrogen fixation, but their widespread effectiveness is constrained by factors such as poor survival and competitiveness of inoculant strains within agricultural soils, as well as variable and context-dependent effects on crop growth (Kaminsky et al., 2019; Batstone et al., 2023). Long-term cropping systems studies have shown that differences in legume use can lead to orders-of-magnitude changes in soil rhizobia abundance. For example, rhizobia abundance was 6.8 × 10⁶ cells g^-1^ dry soil in legume-inclusive rotations compared to 6.1 × 10² cells g^-1^ dry soil in continuous maize (Yan et al., 2014). Therefore, one underexplored solution to increase cash crop productivity may be to select cover crops that enrich soil rhizobial communities for targeted cash crops.

Here, we explore how microbial communities shaped by legume cover crops influence the performance and nodule microbiomes of downstream legume cash crops. In a greenhouse legacy experiment, we inoculated four cash crops (alfalfa, common bean, fava bean, and soybean) with rhizosphere and nodule microbiomes from cover crops under nitrogen-limiting conditions, then assessed plant performance and the resulting nodule bacterial and diazotroph communities via 16S rRNA and *nifH* sequencing, respectively. Using this framework, we address the following questions: 1) How does microbial community composition differ between the rhizosphere and nodules compartments of cover crop species? 2) In the absence of external nitrogen, does inoculum compartment (nodule vs. rhizosphere) or cover crop species influence cash crop growth? 3) Does the taxonomic composition of cover crop inocula predict functional outcomes for cash crop performance? 4) Does prior host filtering in a cover crop inoculum shape cash crops’ ability to form associations with rhizobial symbionts? We hypothesized that cash crop performance is determined by the degree of rhizobial overlap and specificity between the cover crop and cash crop hosts. We found that cash crops differed in host permissiveness, ranging from strict rhizobial specificity to broader recruitment of compatible taxa, and that biomass and nodulation outcomes in a subset of cash crops were shaped by both rhizobial compatibility and additional microbial taxa.

## METHODS

### Phase 1: Cover crop enrichment of microbes and inoculant preparation

In phase 1, we grew eight replicate pots of six legume cover crop species in a mixture of field soil, sand, and calcined clay (Fig. 1). The field soil was collected from the Russell E. Larson Agricultural Research Center (Pennsylvania Furnace, Pennsylvania), where a rotation of corn, soy, wheat, and diverse cover crops has been in place for the past twelve years (Cloutier et al., 2020; Finney et al., 2017). The legume cover crop species included bush clover (*Lespedeza cuneata*), cowpea (*Vigna unguiculata*), crimson clover (*Trifolium incarnatum*), field pea (*Pisum sativum subsp. arvense*), white clover (*Trifolium repens*), and yellow sweet clover (*Melilotus officinalis*). These cover crop species were selected due to their association with a diverse span of rhizobial taxa that form nitrogen-fixing associations with common legume cash crops (Table 1). Ten seeds were planted per pot, and after approximately two weeks of growth, plants were thinned to six to seven per pot. The plants were grown in a randomized block design in the greenhouse and fertilized weekly with nitrogen-free Fahräeus solution (Barker et al., 2006) to encourage nodulation. After 18 weeks of growth, microbial inoculants were prepared from the nodules and rhizosphere of each cover crop replicate.

**Fig. 1.**
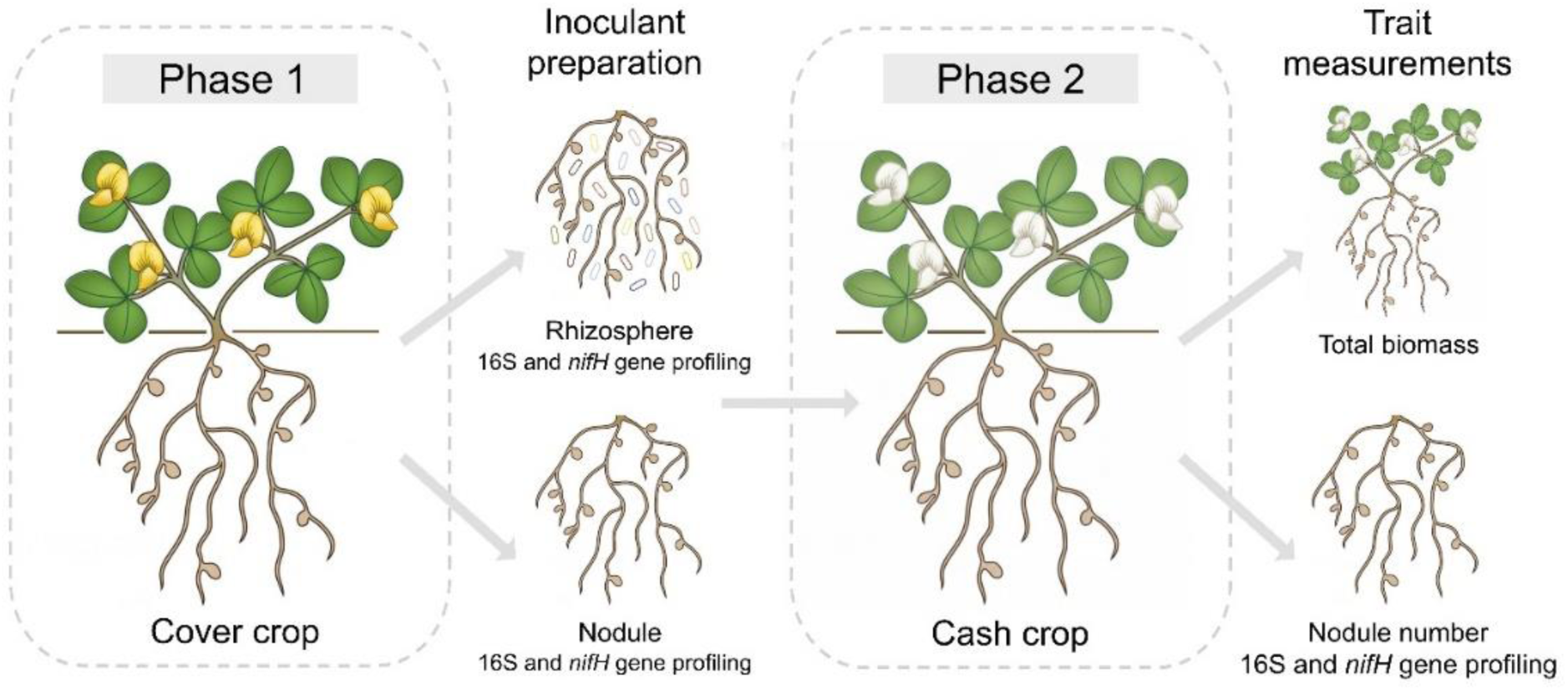
In phase 1, we grew eight pots of six legume cover crop species with six to seven plants per pot (N=48; 6 cover crops, 8 replicates). The legume cover crop species included bush clover, cowpea, crimson clover, field pea, white clover, and yellow sweet clover. After 18 weeks of growth, we assessed the 16S and *nifH* gene profile of the rhizosphere and nodules of each cover crop replicate. In phase 2, we used the rhizosphere and nodules derived from each cover crop species replicate as inocula for four cash crop species that were grown in a randomized block design (N=384; 4 cash crops, 2 compartments, 6 cover crops, 8 replicates). The cash crop species included alfalfa, common bean, fava bean, and soybean. Each block also included one non-inoculated control per cash crop species, which received sterilized DI water instead of inoculum (N=40; 5 cash crops, 8 replicates). To encourage nodulation, all plants received nitrogen-free Fahräeus solution weekly. After 12 weeks of growth, we assessed cash crop performance and the 16S and *nifH* gene profile of cash crop nodules.

**Table 1.**
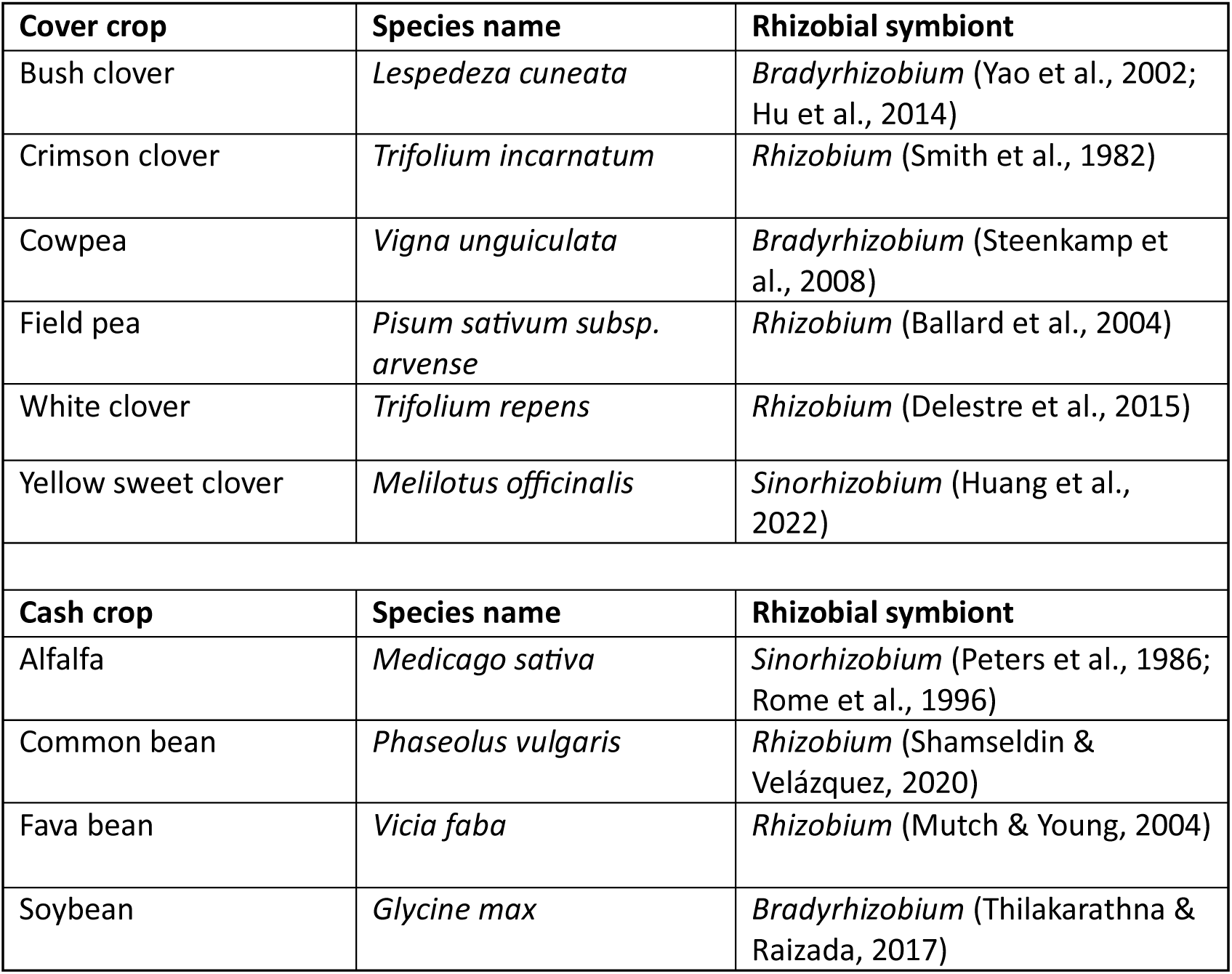
Rhizobial symbionts associated with cover crops and cash crops. The dominant rhizobial genera associated with the six cover crop species and four cash crop species used in phase 1 and phase 2, respectively. Rhizobial symbionts are reported at the genus level to be comparable to 16S and *nifH* results presented here.

To prepare the nodule inoculum, nodules were collected from approximately five plants per pot and placed in a metal-mesh tea strainer. The nodules were surface-sterilized with 15% household bleach for 90 s, followed by rinsing with sterile DI water for 60-90 s. Sterilized nodules were transferred into a tube containing 80 mL of sterile 0.85 % NaCl solution and homogenized for 90–120 s using a tissue homogenizer (Omni International). The homogenate was vortexed and centrifuged at 400 × g for 10 mins at 10°C. The supernatant was collected and centrifuged at 15,000 × g for two mins at 10°C. The final supernatant was discarded, and the pellet was resuspended in 40 mL 0.25× phosphate-buffered saline (PBS) for use as the nodule inoculum. To prepare the rhizosphere inoculum, roots from approximately five plants per pot were collected and placed in a sterile 250 mL media bottle. Sterile DI water was added, and the bottle was shaken for one min to suspend the rhizosphere microbiota. The suspension was centrifuged at 400 × g for 10 mins at 10°C. The resulting supernatant was transferred to a new tube and centrifuged at 15,000 × g for two mins. The supernatant was discarded, and the resulting pellet was resuspended in 40 mL 0.25x PBS to serve as the rhizosphere inoculum.

We used flow cytometry to determine the concentration of the rhizosphere and nodule inoculants. Samples were processed on an Aurora Spectral Analyzer (Cytek Biosciences), which counts cells and records flow rate, allowing us to determine cell counts per mL in each sample. We set a size threshold above 0.2 µm, determined by Flow Cytometry Sub-Micron Particle Size Reference Kit Beads (Invitrogen), to remove small particles and retain microbial cells. Then we removed any plant cells by setting a gate on events that did not show autofluorescence at 488/679 or 561/679 (ex/em), indicative of chlorophyll. We used SYBR green, a DNA probe (Invitrogen, Ex/Em 498/522 nm, final concentration of 0.5 µM), to distinguish cells from other particulate matter. We gated for SYBR-positive events first with a positive control of cultured *Sinorhizobium melioti* (a known nodule endophyte) spiked into soil + SYBR in the 488/525 SYBR and FSC channels. To ensure there was no off-target labeling, we ran a negative control with sterile soil + SYBR and then adjusted our gates to ensure less than 1% false positives. The same gating strategy was used for nodule and rhizosphere samples.

### Phase 2: Cash crop response to nodule and rhizosphere inocula

In phase 2, five cash crop species were grown with the nodule and rhizosphere inocula derived from each cover crop species in phase 1 (Fig. 1). The cash crop species included alfalfa (*Medicago sativa*), common bean (*Phaseolus vulgaris*), fava bean (*Vicia faba*), peanut (*Arachis hypogaea*), and soybean (*Glycine max*). Due to poor nodulation, peanut was excluded from downstream analyses. Prior to planting, the seeds of each cash crop species were surface-sterilized with 15% household bleach for 15 mins, followed by five rinses with sterile DI water. Two surface-sterilized seeds were planted into Deepots (Stuewe & Sons, Inc.) filled with a sterile 1:1 mixture of sand and calcined clay and topped with sterile vermiculite. Pots were arranged in a randomized block design by cash crop species with eight replicates of each cover crop and inoculum compartment in each block (N = 480; 5 cash crops, 2 compartments, 6 cover crops, 8 replicates). One week after planting, the seedlings were inoculated with 6 mL of nodule or rhizosphere inoculum from each cover crop replicate. In addition, each block included one non-inoculated control per cash crop species, which received 6 mL of sterilize DI water instead of inoculum (N = 40; 5 cash crop species, 8 replicates). After two to three weeks of growth, plants were randomly thinned to one plant per pot. To encourage nodulation, all plants received nitrogen-free Fahräeus solution weekly.

After 12 weeks of growth, the proportion of flowered plants was recorded for each cash crop species. Plants were then harvested, and nodules were counted and collected. Following nodule collection, roots and shoots were separated and oven-dried for one week at 60°C prior to biomass measurements. The proportion flowered was analyzed for each cash crop species using logistic regression models for binary data. For total biomass, a value of 0.001 was imputed to all biomass measurements to account for alfalfa samples that were below the scale’s minimum load. Total biomass was evaluated using linear mixed-effects models and two-way ANOVA with Type III sums of squares (lme4 v 2.0-1, Bates et al., 2015; lmerTest v3.2-1, Kuznetsova et al., 2017). Alfalfa total biomass was log-transformed prior to analysis to meet model assumptions. Nodule number was square-root transformed and similarly assessed using linear mixed-effects models with two-way ANOVA with Type III sums of squares. In all models, cover crop, inoculum compartment, and their interaction were included as fixed effects, and block was included as a random intercept to account for variation among experimental blocks. Pairwise contrasts between the nodule and rhizosphere inoculum for each plant trait were performed using Dunnett’s post-hoc procedure with emmeans v2.0.3 (Lenth 2024). To evaluate the effects of inoculum cell count on plant performance, each trait (nodule number and total biomass) was analyzed using a linear mixed-effects model with log-transformed cell counts as a fixed effect and replicate as a random intercept, and then assessed by two-way ANOVAs with Type III sums of squares.

After collection, cash crop nodules were stored for approximately one day at 4°C until processing. Processing followed phase 1 methods for inoculant preparation with slight modifications. Nodules from one to eight cash crop plants per cover crop and inoculum compartment were pooled and placed in a metal mesh tea strainer. Nodules were surface-sterilized with 15% bleach for 30 s, followed by rinsing with sterile DI water for 60 s. Sterilized nodules were transferred into a tube containing approximately 3 mL of autoclaved 0.85 % NaCl solution and homogenized for 90 s using a tissue homogenizer (Omni International). The homogenate was vortexed and centrifuged at 400 × g for 10 mins at 10°C to remove plant debris. The supernatant was then collected and centrifuged at 18,000 × g for five min to produce a bacteria-enriched pellet.

### DNA extraction, 16S rRNA, and nifH amplicon sequencing

In phase 1, nodule and rhizosphere samples were pooled from approximately five plants within each cover crop pot. In total, we collected eight replicate nodule pools and rhizosphere samples per cover crop. In phase 2, nodules from one to eight cash crop plants per cover crop and inoculum compartment were pooled when sufficient nodules were available. Prior to DNA extractions, 1 mL aliquots of the phase 1 nodule and rhizosphere inoculum were pelleted by centrifuging at 15,000 x g for two min. DNA was then extracted from phase 1 and phase 2 nodule pellets using the Qiagen DNeasy Plant Pro Kit (Cat. No. 69206) following manufacturer instructions. Rhizosphere DNA was extracted using the Qiagen DNeasy PowerSoil Kit (Cat. No. 12855), also following manufacturer instructions. The 16S rRNA V4 region was amplified by 515F (Parada et al., 2016) and 806R (Apprill et al., 2015) primers, whereas *nifH* was amplified with the IGK3 (forward) and DVV (reverse) primers (Gaby & Buckley, 2012; Ando et al., 2005). The 16S PCR amplification was performed in 20 µL reactions containing 10 µL Invitrogen Platinum SuperFi PCR Master Mix, 0.4 µL each of forward and reverse primers (10 µM each), 1 µL template DNA, and 8.2 µL PCR-grade water. Thermocycling conditions consisted of an initial denaturation at 98°C for 2 mins, 25 cycles of denaturation at 98°C for 10 s, annealing at 56°C for 20 s, and extension at 72°C for 15 s, followed by a final extension at 72°C for 5 mins. The *nifH* PCR amplification was performed in 20 µL reactions containing 10 µL AmpliTaq Gold 360 Master Mix, 2 µL each of forward and reverse primers (10 µM each), 1.2 µL template DNA, and 4.8 µL PCR-grade water. Thermocycling conditions consisted of an initial denaturation at 95°C for 10 mins, 34 cycles of denaturation at 95°C for 30 s, annealing at 56°C for 45 s, and extension at 72°C for 40 s, followed by a final extension at 72°C for 7 mins. Library preparation and sequencing of 16S and *nifH* amplicons were performed by the Huck Institutes’ Genomics Core Facility on an Illumina NextSeq 2000 to produce 300 bp paired-end reads with each amplicon sequenced in a separate run.

### 16S and nifH sequence data processing

Sequence data were processed using established bioinformatic pipelines (Wagner et al., 2020; Morando et al., 2025). Raw 16S and *nifH* sequences were processed with Cutadapt v3.7 (Martin, 2011) to remove primers. Only read pairs with confirmed primer matches were retained and error rates of 0.1 and 0.15 were used for 16S and *nifH*, respectively, to account for *nifH* primer degeneracy. We examined read quality profiles using FastQC v0.12.1 to assess sequencing quality (Andrews, 2010). For the *nifH* dataset only, we predicted open reading frames (ORFs) from R1 reads using FragGeneScan v1.32 (Rho et al., 2010), then screened ORFs against a nitrogenase Pfam model (PF00142.20; Mistry et al., 2021) using HMMER3 v3.3.2 (Eddy, 2011), retaining only sequences likely to encode nitrogenase reductase. We then used DADA2 v1.30.0 (Callahan et al., 2016) on both datasets to quality-filter and truncate reads, infer amplicon sequence variants (ASVs), and remove chimeras. Truncation lengths were selected based on FastQC quality profiles to retain high-quality bases while ensuring sufficient overlap for merging paired-end reads. Taxonomy was assigned to 16S ASVs using the Ribosomal Database Project (RDP) classifier trained on the RDP training set v. 19 (Cole et al., 2014). ASVs that could not be classified at the kingdom level or were identified as plant sequences were discarded. Taxonomy was assigned to *nifH* ASVs using the nifHdada2 v2.0.5 database (Moynihan & Furbo Reeder, 2023), after which non-target *nifH* homologs were removed to only retain confirmed *nifH* sequences. In addition, *nifH* ASVs were filtered by sequence length to retain only those within the expected amplicon size range. Low-abundance ASVs were removed from both datasets using the kOverA function from the genefilter package v1.92.0 (Gentleman et al., 2023) based on minimum count and prevalence thresholds. This retained 98% and 95% of original reads for the 16S and *nifH* datasets, respectively. Next, samples with fewer than 10,000 reads were removed, resulting in the exclusion of two samples from the *nifH* dataset but none from the 16S dataset. The final 16S dataset contained 35,367,620 reads across 135 samples with a mean (±SD) library size of 260,056 ± 76,040 reads per sample and 5,128 ASVs, while the final *nifH* dataset contained 10,505,380 reads across 133 samples with a mean (±SD) library size of 78,988 ± 48,302 reads per sample and 254 ASVs. Finally, we applied a centered log-ratio (CLR) transformation to each count dataset using the ALDEx2 R package v1.42.0 (Fernandes et al., 2013) to account for the compositional nature of microbiome data (Gloor et al., 2017).

### Microbiome analyses

All data analyses were performed in R using RStudio version 4.3.2 with data wrangling and visualization conducted using the tidyverse suite of packages v2.0.0 (Wickham et al., 2019). Alpha diversity metrics (Richness and Shannon diversity indices) were calculated using the phyloseq R package v1.54.2 (McMurdie & Holmes, 2013). These metrics were modeled using the lme4 R package v2.0-1 (Bates et al., 2015) with compartment, cover crop, and their interaction, as well as sequencing depth, as fixed predictors, and block as a random-intercept term. We assessed these linear mixed-effect models using ANCOVA, then adjusted the p-values for multiple comparisons (Benjamini & Hochberg, 1995). A canonical analysis of principal components (CAP) ordination using Aitchison distance was performed on the CLR-transformed bacterial counts, constrained by compartment, cover crop, and their interaction, with sequencing depth partialled out to remove noise. Separate models were run for each compartment to test the effects of cover crop identity with sequencing depth partialled out. We then used an ANOVA-like permutation test for Constrained Correspondence Analysis using the vegan R package v2.7-3 (Oksanen et al., 2022) to determine whether the bacterial taxa counts were explained by the constraining predictors.

To assess whether the composition of cover crop inoculum communities predicted cash crop biomass and nodule number, we performed a redundancy analysis (RDA) on CLR-transformed ASV abundances constrained by each plant trait, with log-transformed sequencing depth partialled out. Separate RDA models were fit for each cash crop, plant trait, and microbial dataset (bacterial and diazotroph), and model significance was assessed using an ANOVA-like permutation test. For significant models, taxon loadings were extracted and the direction of the biplot arrow for the constraining trait was used to determine the association between taxon loadings and plant performance. Candidate taxa were prioritized based on RDA1 loading magnitude, prevalence across samples, and biological relevance. For each prioritized taxon, a linear mixed-effects model was fit with the relevant plant trait as the response, taxon CLR abundance, inoculum compartment, and their interaction as fixed effects, log-transformed sequencing depth as an additional fixed effect, and cover crop identity as a random effect. Follow-up models were fit separately for nodule and rhizosphere inoculum, retaining taxon CLR abundance and log-transformed sequencing depth as fixed effects and cover crop identity as a random effect. Significance of all models was assessed using ANOVA with Type III sums of squares implemented in the lmerTest package v3.2-1 (Kuznetsova et al., 2017). P-values were adjusted using the Benjamini-Hochberg procedure separately for the bacterial and diazotroph dataset (Benjamini & Hochberg, 1995).

To assess the degree to which cash crop nodule communities reflected the composition of their corresponding cover crop inoculum, we calculated a shared read proportion for each cash crop nodule sample using the *nifH* sequence data. For each cash crop nodule sample, we identified the set of ASVs detected in the corresponding cover crop source sample, matched by cover crop identity and inoculum compartment (nodule or rhizosphere), and calculated the proportion of total reads in the cash crop nodule sample that were shared with that source. Values close to 1 indicate a high degree of overlap between the cash crop nodule community and the corresponding cover crop inoculum, while values close to 0 indicate low overlap between the two.

## RESULTS

### Cover crop identity shapes bacterial and diazotroph community composition in rhizosphere and nodule microbiomes

In phase 1, we examined how microbial community composition varied by species identity and compartment across six legume cover crops (Fig. 1; Table 1). We characterized both the broader bacterial community and the diazotroph community—the nitrogen-fixing bacteria commonly associated with legumes—using 16S rRNA and *nifH* marker gene sequencing, respectively. A permutational ANOVA confirmed significant effects of cover crop identity (F_5,82_=4.5, p<0.001), compartment (F_1,82_=21, p<0.001), and their interaction (F_5,82_=2.3, p<0.001) on bacterial community composition (Fig. 2A), with significant effects of cover crop identity observed in both nodule (F_5,41_=2.6, p<0.001) and rhizosphere (F_5,40_=3.4, p<0.001) compartments. These patterns were mirrored in the diazotroph community, in which cover crop identity (F_5,80_=5.0, p<0.001), compartment (F_1,80_=2.8, p<0.001), and their interaction (F_5,80_=2.0, p<0.001) all significantly influenced *nifH* community composition (Fig. 2B), with significant effects of cover crop identity in both nodule (F_5,41_=5.3, p<0.001) and rhizosphere (F_5,38_=2.5, p<0.001) compartments. Rhizosphere communities consistently exhibited higher Shannon diversity than nodules for both 16S (Fig. 2C; Supplemental Table 1) and *nifH* (Fig. 2D; Supplemental Table 2) amplicons. Nodule microbiomes were dominated by the same limited number of genera in both bacterial and diazotroph communities (Supplemental Fig. 1; Supplemental Fig. 2), though a greater number of rhizobial symbiont ASVs were detected by *nifH* than 16S sequencing (Supplemental Table 3). Dominant genera varied by cover crop: *Rhizobium* was most abundant in crimson clover, field pea, and white clover; *Sinorhizobium* in yellow sweet clover; and *Bradyrhizobium* in bush clover and cowpea. These results indicate that bacterial and diazotroph community composition varied significantly with cover crop identity in both nodules and rhizosphere, while diversity measures were consistently higher in the rhizosphere across all cover crops.

**Fig. 2.**
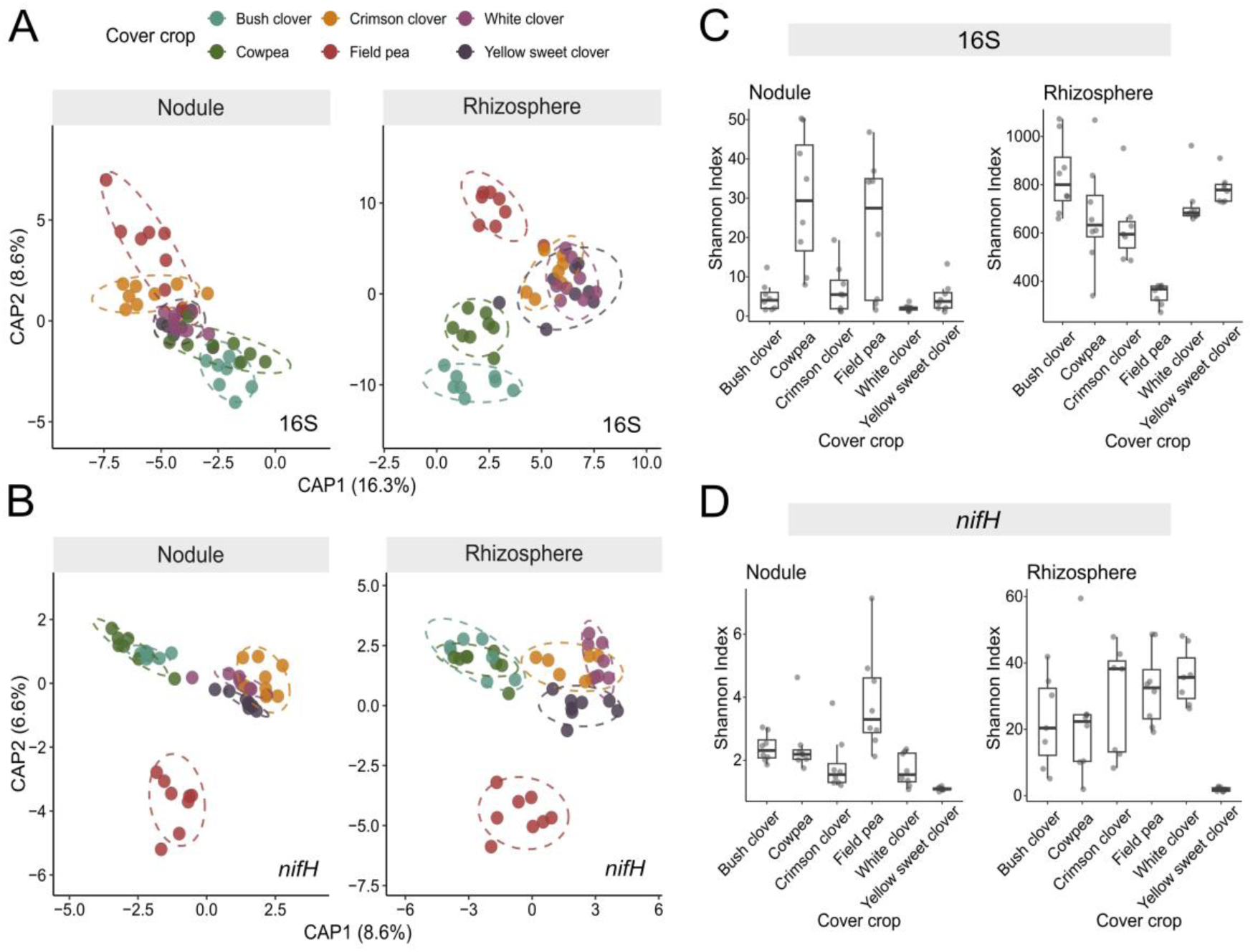
Characterization of cover crop nodule and rhizosphere bacterial and diazotroph communities. Constrained analysis of principal coordinates (CAP) ordination of (A) bacterial and (B) diazotroph communities constrained by cover crop species and compartment (nodule vs. rhizosphere). Bacterial community composition was significantly influenced by cover crop (F_5,82_=4.5, p<0.001), compartment (F_1,82_=21, p<0.001), and their interaction (F_5,82_=2.3, p<0.001). Diazotroph community composition similarly was significantly influenced by cover crop (F_5,80_=5.0, p<0.001), compartment (F_1,80_=2.8, p<0.001), and their interaction (F_5,80_=2.0, p<0.001). Shannon diversity of (C) bacterial and (D) diazotroph communities in nodules and rhizospheres across cover crop species. Bacterial Shannon diversity was significantly influenced by cover crop (F_5,76_=12.2, p<0.001), compartment (F_1,78_=947.6, p<0.001), and their interaction (F_5,77_=13.4, p<0.001). Diazotroph Shannon diversity was significantly influenced by cover crop (F_5,74_=3.8, p<0.05), compartment (F_1,79_=15.6, p<0.001), and their interaction (F_5,74_=5.0, p<0.01). P-values for Shannon diversity were adjusted using the Benjamini-Hochberg method.

### Cash crop performance varied among species in response to cover crop inoculum

In phase 2, we assessed how inocula from nodule and rhizosphere communities shaped by cover crops influenced cash crop performance (Fig. 1). After 12 weeks of growth, we quantified the number of nodules, biomass accumulation, and proportion of flowered plants for each cash crop (alfalfa, common bean, fava bean, and soybean). Across cash crops, the effects of inoculum compartment (rhizosphere vs. nodule) and cover crop identity on plant performance varied considerably. Inoculum compartment had no measurable effect on nodule number for any cash crop (Fig. 3A; Supplemental Table 4), whereas cover crop identity significantly influenced nodule number in all cash crop species, with particularly large effects in alfalfa (F₅,₅₉=12.1, p<0.001). A significant interaction between inoculum compartment and cover crop identity occurred only in common bean (F₅,₈₄=2.9, p=0.019), whereby bush clover, cowpea, and field pea rhizosphere inocula resulted in greater nodule number, and crimson clover, white clover, and yellow sweet clover nodule inocula resulted in greater nodule number. For total biomass, inoculum compartment effects were similarly limited with a substantial positive effect only in common bean (F₁,₇₇=19.3, p<0.001), which was driven by an increase in biomass when grown with cover crop rhizosphere inoculum (Fig. 3B; Supplemental Table 5). In contrast to nodule number, cover crop identity effects on total biomass were detected only in alfalfa (F₅,₆₃=7.6, p<0.001) and fava bean (F₅,₇₂=2.39, p=0.046), suggesting a direct relationship between increased nodulation and plant growth for these two cash crops. No interactions were observed for total biomass, indicating that biomass responses to inoculum and cover crop identity were largely independent. Cover crop identity did not influence the proportion of plants that flowered across cash crops (Supplemental Fig. 3; Supplemental Table 6). In contrast, inoculum compartment significantly affected flowering in common bean (X^2^_1_=8.29, p=0.004), with a higher proportion of plants flowering when receiving rhizosphere inoculum compared to nodule inoculum for all cover crops except cowpea. Inoculum cell count had no significant effects on nodule number or total biomass for any of the cash crops (Supplemental Table 7). Overall, cover crop identity was the primary determinant of cash crop response to inoculation, driving nodulation across all species and biomass accumulation in alfalfa and fava bean specifically. Inoculum compartment had a comparatively small effect, evident only in common bean.

**Fig. 3.**
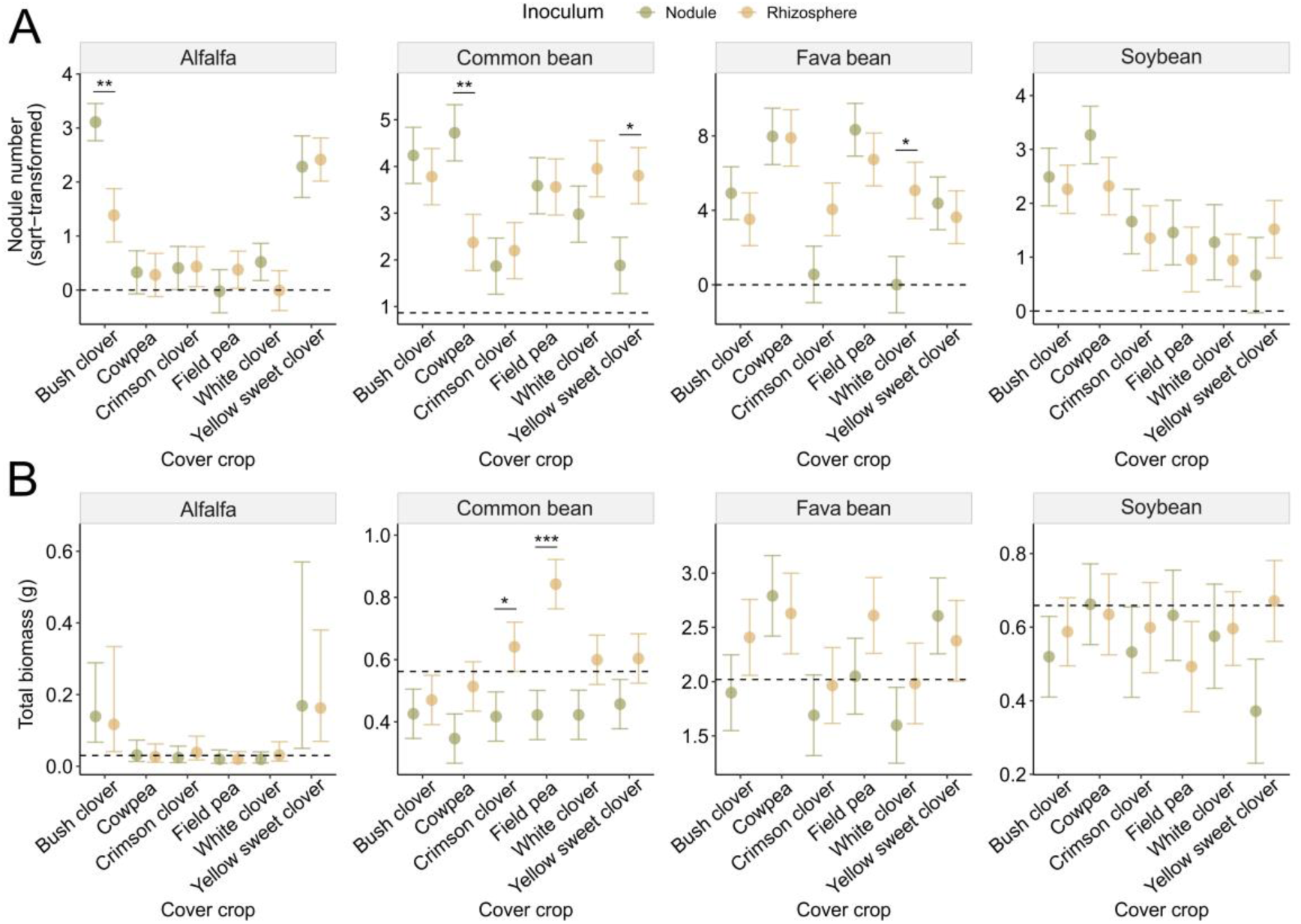
Effect of cover crop inocula on cash crop traits in phase 2. (A) Square-root transformed nodule number for each cash crop grown in nodule or rhizosphere inoculum from each cover crop and (B) Total biomass for each cash crop grown in nodule or rhizosphere inoculum from each cover crop. Error bars show the standard error for the estimated marginal mean values and dashed lines indicate median nodule number or total biomass for non-inoculated control. Pairwise contrasts between the nodule and rhizosphere inoculum for nodule number and total biomass were performed using Dunnett’s post-hoc procedure (* = p < 0.05, ** = p < 0.01, *** = p < 0.001).

### Taxonomic composition of cover crop rhizosphere inoculum predicts common bean biomass and soybean nodulation success

In phase 2, we assessed whether taxa within cover crop inoculum communities were associated with cash crop biomass accumulation and nodulation success. Cover crop inoculum community composition predicted cash crop performance in a species-specific manner, with significant associations detected for common bean biomass and soybean nodule number but not for alfalfa or fava bean. Redundancy analysis (RDA) showed that bacterial community composition explained more variance in common bean biomass (Adj. R²=0.023, p=0.001) than diazotroph community composition (Adj. R²=0.007, p=0.011). Taxa prioritized from these RDA models were then evaluated individually within rhizosphere and nodule inoculum, with several taxa showing significant associations with biomass in rhizosphere inoculum, but not nodule inoculum (Figure 4A). CLR-transformed abundance of *Novosphingobium* ASV 21 (*nifH*) was marginally positively associated with common bean biomass (F_1,38_=5.2, adj. p=0.056), as well as *Rhizobium* ASV 10 (16S; F_1,90_=6.2, adj. p=0.035) and *Sphingomonas* ASV 18 (16S; F_1,90_=14.4, adj. p=0.003). In contrast, abundance of *Asaia* ASV 84 (*nifH*) was negatively associated with biomass (F_1,42_=7.5, adj. p=0.036), along with *Methylotenera* ASV 199 (16S; F_1,90_=9.3, adj. p=0.015) and *Rubrivivax* ASV 167 (16S; F_1,90_=8.9, adj. p=0.003). While bacterial community composition showed no effect on soybean nodule number, diazotroph inoculum community composition was a significant predictor (Adj. R²=0.020, p=0.001). At the taxon level, associations were specific to rhizosphere inoculum (Figure 4B): *Bradyrhizobium* ASV 3 (*nifH*) abundance was positively associated with nodule number (F_1,27_=10.6, adj. p=0.012), while *Rhizobium* ASV 5 (*nifH*) abundance was negatively associated with nodule number (F_1,27_=7, adj. p=0.027). These results indicate that both bacterial and diazotrophic taxa in cover crop rhizosphere inoculum are associated with cash crop biomass accumulation and nodulation, with the direction of the relationship varying by taxon.

**Fig. 4.**
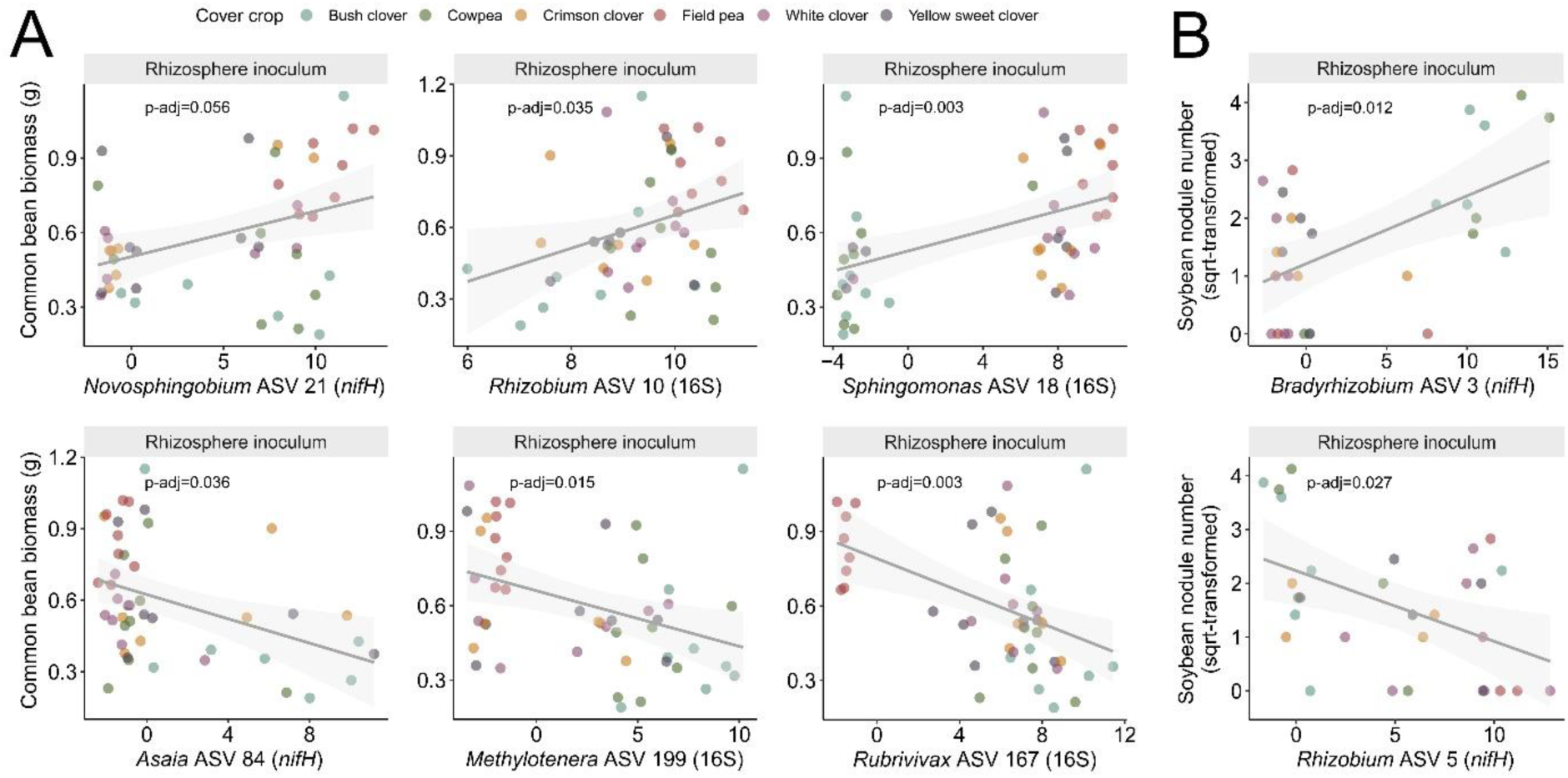
Bacterial and diazotroph taxa associated with common bean biomass accumulation and soybean nodulation in cover crop rhizosphere inoculum. Taxa were prioritized from RDA models constrained by the corresponding plant trait and were significant positive or negative predictors of that trait in rhizosphere-specific linear mixed-effects models. (A) CLR-transformed abundance of bacterial and diazotrophic taxa versus common bean total biomass. (B) CLR-transformed abundance of diazotrophic taxa versus square root-transformed soybean nodule number. Points represent individual samples colored by cover crop identity; lines show fitted linear trends with 95% confidence intervals (shaded). Dataset is indicated in parentheses (16S: bacterial; nifH: diazotroph); p-values were adjusted within each dataset using the Benjamini-Hochberg procedure.

### Cash crop nodule diazotroph communities reflect species-specific selection from cover crop inoculum

To assess the influence of cover crop inoculum on cash crop nodule microbiomes, we characterized the microbial community composition of nodules pooled across replicate plants. However, some cover crop–inoculum combinations yielded an insufficient number of cash crop nodules for microbiome assessment. Cash crop nodules were largely dominated by their previously described rhizobial symbionts across cover crops for each inoculum compartment (nodule or rhizosphere) in bacterial (Supplemental Fig. 4; Supplemental Table 8) and diazotroph communities (Table 1; Fig. 5A; Supplemental Table 9). However, non-rhizobial bacterial genera were detected in nodules of all cover crop species (Supplemental Fig. 4). Notably, *Pseudomonas* was abundant across nodule bacterial communities (16S) in all cash crops, with the highest relative abundance observed in soybean nodules. Within the *nifH* dataset, non-*Rhizobium* rhizobial genera (i.e., *Bradyrhizobium* and *Sinorhizobium*) constituted a substantial proportion of the common bean nodule diazotroph community when receiving cover crop nodule inoculum (Fig. 5A; Supplemental Table 9). These genera were enriched by cover crop species that associate with them, suggesting common bean is a relatively permissive host for diverse rhizobial taxa when present in high abundance, such as in the nodule environment (Supplemental Fig. 5). Common bean’s proportion of shared *nifH* reads with cover crop inoculum was also lower and more variable, particularly for nodule-derived inoculum, consistent with this permissiveness (Fig. 5B). Alfalfa and soybean showed consistently high overlap with cover crop inoculum (near 1.0) regardless of inoculum compartment. Fava bean showed the lowest and most variable overlap of the four cash crops, in both nodule- and rhizosphere-derived inoculum. These host-specific patterns were also reflected at the *nifH* ASV level: common bean and fava bean nodules were each composed of numerous *Rhizobium* ASVs. In common bean, ASV 5 typically dominated regardless of cover crop identity or inoculum compartment, whereas fava bean’s dominant ASV varied by both (Fig. 5C). In contrast, a single ASV (ASV 1) accounted for the overwhelming majority of *Sinorhizobium* relative abundance in alfalfa nodules, whereas soybean nodules were similarly dominated by a small set of *Bradyrhizobium* ASVs (ASV 3 and ASV 7), though their relative dominance varied by inoculum compartment, indicating more restrictive selection for these cash crops. Within each cover crop, the dominant *nifH* ASV was generally consistent between nodule and rhizosphere compartments, at similar relative abundance, with the exception of field pea (Supplemental Fig. 6). This suggests that cover crop compartment did not majorly affect symbiont ASV availability for cash crop selection. Together, these results suggest that cash crop hosts differ in how selectively they assemble their nodule diazotroph communities from available cover crop inoculum.

**Fig. 5.**
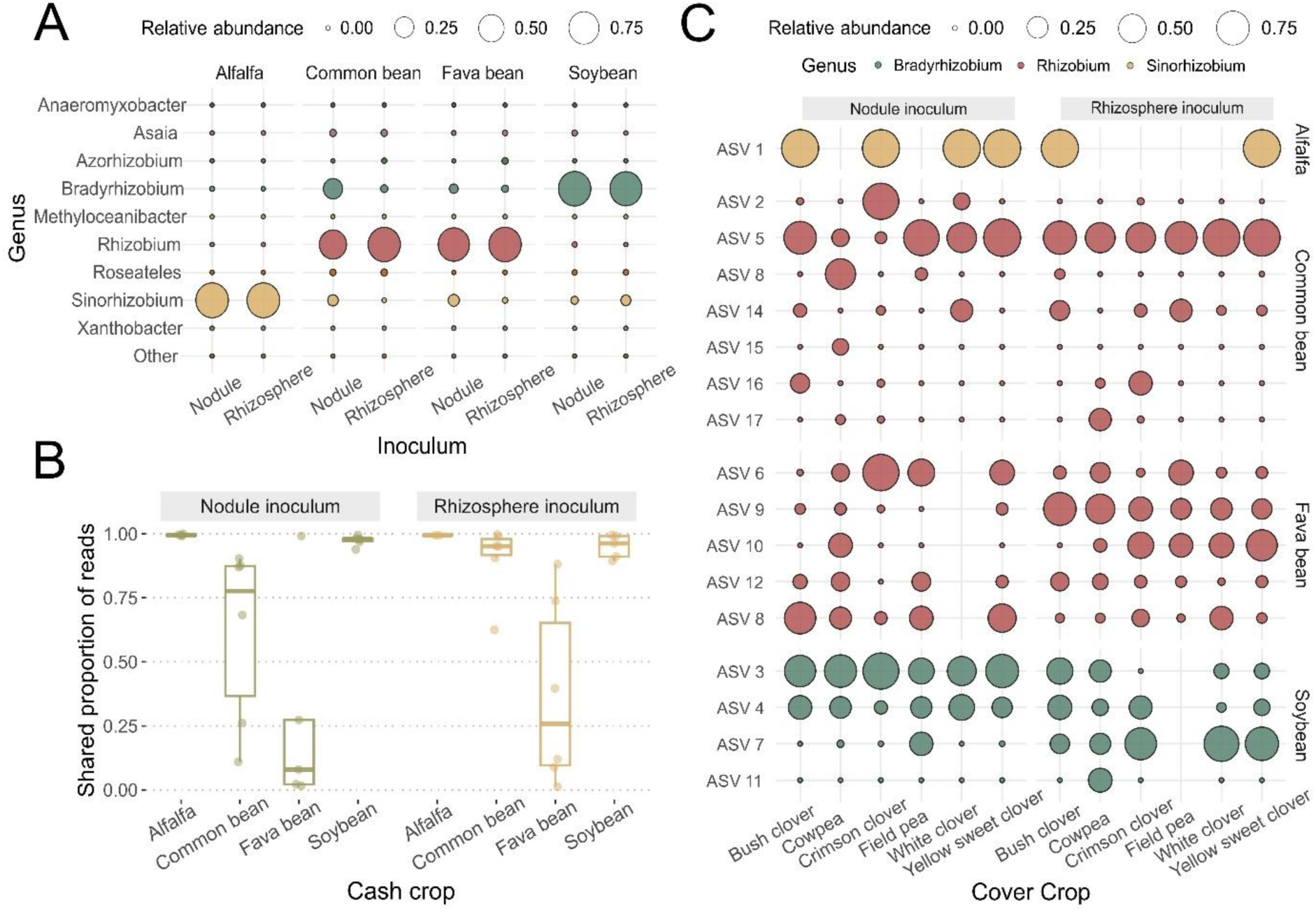
Composition and selectivity of cash crop nodule diazotroph communities. (A) Genus-level composition of diazotroph (*nifH*) communities in cash crop nodules, shown across cover crop inoculum compartment (nodule or rhizosphere). Bubble size reflects mean relative abundance of each genus within each cash crop × inoculum compartment combination, averaged across cover crops. Low-abundance genera outside the top taxa shown are grouped as “Other.” (B) Proportion of *nifH* reads in cash crop nodules belonging to taxa that were also detected in the corresponding cover crop inoculum, reflecting the degree of overlap between the cash crop nodule community and the cover crop inoculum it received. Each point corresponds to a specific cover crop that supplied the inoculum, faceted by inoculum compartment (nodule or rhizosphere). (C) ASV-level composition of the *nifH* rhizobial symbiont genus associated with each cash crop (*Sinorhizobium* for alfalfa; *Rhizobium* for common bean and fava bean; *Bradyrhizobium* for soybean) within cash crop nodules. For each cash crop, the minimal set of ASVs collectively accounting for >97% of mean relative abundance across samples is shown. Bubble size reflects relative abundance within each cover crop × inoculum compartment combination; color indicates genus. No bubble indicates that no nodules formed, preventing analysis of cash crop nodule microbiomes.

## DISCUSSION

Understanding how legume–rhizobium compatibility shapes symbiotic nitrogen fixation is critical for predicting microbial contributions to agricultural productivity (Herridge et al., 2008; Gopalakrishnan et al., 2015). Here, we examined how microbial communities shaped by legume cover crops influenced nodule microbial communities and plant performance in subsequent legume cash crops. We identified taxa in cover crop inocula that predicted downstream symbiotic performance. In particular, common bean biomass accumulation was predicted by both rhizobial and non-rhizobial taxa, while soybean nodulation success was predicted by the presence of its compatible rhizobial symbiont. These results reveal a previously underexplored role of microbial compatibility between cover crops and cash crops, extending beyond rhizobial symbionts, in shaping downstream symbiotic outcomes for cash crops. Bacterial and diazotroph diversity was much higher in rhizosphere communities than in nodule communities of cover crops, reflecting strong host filtering during cover crop nodule formation. This same selectivity carried into the subsequent cash crops, as most hosts recruited a consistent set of rhizobial symbionts into their nodules regardless of cover crop inoculum, with only slight variation in relative abundance and diversity, particularly for common bean. Despite this consistency in symbiont identity, cover crop identity still significantly affected cash crop nodulation and, to a lesser extent, total biomass, suggesting legume cover crops can leave lasting effects on subsequent cash crop performance.

The strong influence of cover crop identity on both rhizosphere and nodule community composition suggests that host-specific root traits and/or exudate chemistry, rather than compartment alone, are primary drivers of microbial community assembly. Legumes rely on root exudates, including flavonoids and other signaling compounds, to recruit compatible rhizobia and establish symbiotic partnerships (Maxwell et al., 1989; Walker et al., 2020). This exudate-driven recruitment is highly species specific, with exudation patterns varying even among cultivars of the same species (Seitz et al., 2023). This compatibility is mediated in part by Nod factor signaling: host-derived flavonoids induce rhizobial Nod factor production, and rhizobia secrete these Nod factors to trigger the early symbiotic responses needed for nodule formation (Kato et al., 2026). Consistent with this signaling-based compatibility, cover crop nodules in our dataset were dominated by known rhizobial symbionts characteristic of each host, indicating that host specificity constrains rhizobial establishment even when a diverse pool of taxa is available in the surrounding soil. This same Nod factor signaling also modulates root exudate profiles and shapes root and rhizosphere microbiota assembly (Tao et al., 2024), which may help explain why rhizosphere communities were similarly structured by cover crop identity here. Other host traits, such as additional exudate compounds or root architecture, likely also contribute to these patterns (Kato et al., 2026; Stewart et al., 2026). These results suggest that cover crops generate distinct microbial legacies in the nodules and rhizosphere, which may in turn contribute to the performance of subsequent cash crops.

Among the cash crops, alfalfa and fava bean showed responses in both biomass accumulation and nodulation to cover crop identity, although fava bean responses were highly variable across cover crop inocula. In alfalfa, this response was particularly pronounced when cover crop inocula contained compatible *Sinorhizobium* symbionts, consistent with its narrow reliance on *S. meliloti* (or *S. medicae* in some cases) for effective symbiotic nitrogen fixation (Bromfield & Cloutier, 2026; Carelli et al., 2000; Silva et al., 2007), highlighting the importance of compatible symbiont identity for alfalfa performance. In common bean and soybean, cover crop identity affected nodulation but did not consistently translate into differences in biomass, suggesting that nodule number alone is an imperfect predictor of plant performance for these species. This disconnect may reflect variation in rhizobial effectiveness, as some strains can form nodules without providing substantial benefits to their hosts (Thrall et al., 2011; Pahua et al., 2018). Thus, differences in the rhizobial communities introduced with cover crop inocula may have affected not only nodule formation but also the benefits provided to the host following symbiosis. Such variation in symbiotic outcomes is consistent with host control of the legume– rhizobia symbiosis, wherein legumes use pre-infection partner choice and post-infection sanctions to preferentially support more beneficial rhizobia (Porter et al., 2024). Although rhizosphere and nodule communities differed in composition and diversity within cover crop hosts, these differences generally did not correspond to differences in cash crop growth, suggesting that compartment-specific taxa were not consistently linked to plant performance. Common bean was the exception: biomass and the proportion of flowering plants were greater under rhizosphere inoculation than under nodule inoculation, indicating that the inoculum compartment, in addition to cover crop identity, can shape plant performance and implicating non-rhizobial members of the rhizosphere community in promoting common bean growth. Together, these results suggest that inoculum composition can influence plant performance at both ends of the host specificity spectrum, but through contrasting mechanisms.

Taxonomic composition of cover crop inocula predicted cash crop performance for only two of the four cash crops tested, and specific taxa within these communities were associated with host performance outcomes. In common bean, *Rhizobium* was the strongest positive predictor of biomass accumulation, consistent with its established role as the primary rhizobial symbiont for this species (Shamseldin & Velázquez, 2020). Two other positive predictors, *Novosphingobium* and *Sphingomonas*, are frequently associated with the common bean rhizosphere (Stopnisek & Shade, 2021) and have been found colonizing common bean roots directly (Barraza et al., 2020), suggesting a consistent association with this host. In particular, *Novosphingobium* can produce the phytohormone indole acetic acid (Aserse et al., 2013) and fix nitrogen in association with a range of plant species (Huang et al., 2024; Rangjaroen et al., 2015), offering a plausible route by which it could support common bean growth independently of, or in combination with, rhizobial nitrogen fixation. By contrast, *Methylotenera*, *Rubrivivax*, and *Asaia* were negative predictors of biomass, though their ecological roles in the common bean rhizosphere remain poorly characterized. A similar pattern of taxon-specific prediction emerged in soybean, though through a different mechanism. *Bradyrhizobium* abundance in the rhizosphere of *Bradyrhizobium*-associated cover crops predicted greater soybean nodulation, whereas *Rhizobium* abundance in the rhizosphere of *Rhizobium*-associated cover crops predicted reduced nodulation. This opposing pattern suggests that rhizobial communities associated with different cover crop hosts may have distinct effects on subsequent soybean–rhizobia interactions. Overall, these results point to specific taxa, several with plausible symbiotic or growth-promoting functions, as the primary drivers of cash crop performance, rather than broad shifts in rhizosphere community composition.

Diazotroph communities within cash crop nodules differed by host species, indicating that cash crops actively filter and select compatible symbionts from the available microbial pool rather than simply reflecting inoculum composition. Notably, this filtering was largely consistent whether the inoculum was the diverse rhizosphere community or the lower-diversity, cover crop-filtered nodule community, though compartment had a modest effect in some cash crops. Host species ranged from generalist to specialist in this filtering: common bean and fava bean permitted a broader range of rhizobial taxa into nodules, while alfalfa and soybean restricted nodule occupancy to a narrower range of rhizobial taxa. However, host selectivity did not always translate into occupancy by the expected primary symbiont. Common bean, a notoriously promiscuous host (Efstathiadou et al., 2021; Shamseldin & Velázquez, 2020), did not primarily associate with *Rhizobium*, instead hosting non-*Rhizobium* rhizobial taxa or *Flavobacterium* depending on the inoculum compartment and microbial community profiled. *Flavobacterium* has been detected in nodules of several legume species, including common bean, though how host traits shape its colonization remains unclear (Rocha et al., 2023; Yu et al., 2025). Even soybean, a more selective host, showed substantial nodule colonization by *Pseudomonas* in the bacterial community rather than its primary symbiont *Bradyrhizobium*. *Pseudomonas* co-inoculation with *Bradyrhizobium* can have contrasting effects on soybean nodule number and acetylene reduction activity, with one strain increasing and another decreasing these traits (Chebotar et al., 2001). However, given that cover crop inocula had only marginal effects on soybean biomass relative to the non-inoculated control, *Pseudomonas* enrichment may have limited *Bradyrhizobium’s* functional contribution despite soybean’s overall selectivity. This aligns with broader evidence that non-rhizobial nodule occupants can modulate symbiosis outcomes, either enhancing or disrupting plant-rhizobia interactions (Martínez-Hidalgo & Hirsch, 2017; Kosmopoulos et al., 2024). Finally, the differences in host permissiveness observed here may also explain why rhizosphere composition was more predictive of performance in some cash crops than others: generalist hosts offer more opportunity for inoculum taxa to influence colonization and outcomes, while selective hosts constrain this regardless of what’s available.

Together, our results show that cover crop microbial legacies shape cash crop performance outcomes through multiple host-specific mechanisms. Selective hosts such as soybean and alfalfa depended on the presence of compatible rhizobial partners within the cover crop inoculum, while a more permissive host like common bean benefited from a broader mix of rhizobial and non-rhizobial taxa. These findings suggest that cover crop selection could be tailored to cash crop host range, matching specific symbionts for more selective hosts and leveraging broader microbial diversity for more permissive ones. However, these findings are based on controlled greenhouse conditions, and additional work is needed to test whether they hold up under more variable, field conditions. Further, extending the duration of cover crop conditioning or monitoring microbial succession across multiple growing seasons could provide deeper insight into how these microbial legacies develop and persist. Similarly, refining cover crop–cash crop pairings based on strain-level rhizobial compatibility, while avoiding cover crops that enrich rhizobia incompatible with the cash crop, may reveal stronger and more consistent benefits for cash crop nodulation and nitrogen fixation. Additional mechanisms may also influence cash crop performance following prior cover crop cultivation and warrant further investigation, including shifts in root exudate composition, changes in microbial community functions, and altered nutrient availability (Seitz et al., 2024; Ghosh et al., 2024). Overall, these findings highlight the potential to strategically select cover crops that foster effective microbial symbionts in subsequent legumes, improving both crop productivity and sustainable nitrogen management in legume-based systems.

## Authorship credits

L.T.B. conceived the research and designed the experiments. E.L.P. and E.B. executed the phase 1 experiment. E.L.P. and J.E.H. prepared and assessed the microbial inoculants. E.L.P., E.B., A.P.J., A.L.S., and K.M. executed the phase 2 experiment. C.L.D., A.L.S., and K.M.C. processed samples and performed DNA extractions. K.M.C. performed the analyses and drafted the manuscript. All the authors reviewed, edited, and approved the final paper.

## Acknowledgments

We would like to acknowledge the Huck Institutes’ Flow Cytometry Core Facility (RRID:SCR_024460) for use of the BD Fortessa Flow Cytometer, the Penn State Institute for Computational and Data Sciences (RRID:SCR_025154) for providing access to computational infrastructure through the Roar Core Facility (RRID:SCR_026424), and the Huck Institutes’ Genomics Core Facility (RRID:SCR_023645) for library preparation and use of the Illumina NextSeq 2000. We would also like to thank Maria Alejandra Gil-Polo, Sohini Guha, and Patrick Sydow for their help picking nodules, Jason Kaye and Brosi Bradley for the long-term cover crop experiment soil used in phase 1, and Scott Diloreto for greenhouse support and plant care advice. This work is/was supported [in part] by Hatch Appropriations under Project #PEN04760 and Accession #1025611, and by the U.S. Department of Agriculture, National Institute of Food and Agriculture under Award #2022-67013-36860 to LTB. While working on this project, JEH was funded by a NIFA Predoctoral Fellowship Award #2023-67011-40517, and KCM was funded by a National Science Foundation Graduate Fellowship. Any opinions, findings, conclusions, or recommendations expressed in this publication are those of the author(s) and do not necessarily reflect the view of the U.S. Department of Agriculture.

## REFERENCES

Ando, S., Goto, M., Meunchang, S., Thongra-ar, P., Fujiwara, T., Hayashi, H., & Yoneyama, T. (2005). Detection of nifH Sequences in Sugarcane (*Saccharum officinarum* L.) and Pineapple (*Ananas comosus* [L.] Merr.). Soil Science and Plant Nutrition, 51(2), 303–308. 10.1111/j.1747-0765.2005.tb00034.x

Andrews, S. (2010). FastQC: a quality control tool for high throughput sequence data. Available online: http://www.bioinformatics.babraham.ac.uk/projects/fastqc

Apprill, A., McNally, S., Parsons, R., & Weber, L. (2015). Minor revision to V4 region SSU rRNA 806R gene primer greatly increases detection of SAR11 bacterioplankton. Aquatic Microbial Ecology, 75(2), 129–137. 10.3354/ame01753

Aserse, A. A., Räsänen, L. A., Aseffa, F., Hailemariam, A., & Lindström, K. (2013). Diversity of sporadic symbionts and nonsymbiotic endophytic bacteria isolated from nodules of woody, shrub, and food legumes in Ethiopia. Applied Microbiology and Biotechnology, 97(23), 10117– 10134. 10.1007/s00253-013-5248-4

Ballard, R. A., Charman, N., McInnes, A., & Davidson, J. A. (2004). Size, symbiotic effectiveness and genetic diversity of field pea rhizobia (*Rhizobium leguminosarum* bv. *Viciae*) populations in South Australian soils. Soil Biology and Biochemistry, Nitrogen Fixation in Australian Agricultural Systems: 13th Australian Nitrogen Fixation Conference, 36(8), 1347–1355. 10.1016/j.soilbio.2004.04.016

Barker, D. G., Pfaff, T., Moreau, D., Groves, E., Ruffel, S., Lepetit, M., Whitehand, S., Maillet, F., Nair, R.M., Journet, E.P. (2006). Growing *M. truncatula*: choice of substrates and growth conditions. The Medicago truncatula handbook. 1–28.

Barraza, A., Vizuet-de-Rueda, J. C., & Alvarez-Venegas, R. (2020). Highly diverse root endophyte bacterial community is driven by growth substrate and is plant genotype-independent in common bean (*Phaseolus vulgaris* L.). PeerJ, 8, e9423. 10.7717/peerj.9423

Bates, D., Mächler, M., Bolker, B., & Walker, S. (2015). Fitting Linear Mixed-Effects Models Using lme4. Journal of Statistical Software, 67, 1–48. 10.18637/jss.v067.i01

Batstone, R. T., Ibrahim, A., & MacLean, L. T. (2023). Microbiomes: Getting to the root of the rhizobial competition problem in agriculture. Current Biology, 33(14), R777–R780. 10.1016/j.cub.2023.05.053

Benjamini, Y., & Hochberg, Y. (1995). Controlling the False Discovery Rate: A Practical and Powerful Approach to Multiple Testing. Journal of the Royal Statistical Society. Series B (Methodological), 57(1), 289–300.

Bromfield, E. S. P., & Cloutier, S. (2026). Complete genome sequences of two effective nitrogen-fixing *Sinorhizobium medicae* strains isolated from *Medicago sativa* and *Melilotus albus* in Canada. Microbiology Resource Announcements, 15(5), e00105–26. 10.1128/mra.00105-26

Burghardt, L. T. (2020). Evolving together, evolving apart: Measuring the fitness of rhizobial bacteria in and out of symbiosis with leguminous plants. New Phytologist, 228(1), 28–34. 10.1111/nph.16045

Burghardt, L. T., & diCenzo, G. C. (2023). The evolutionary ecology of rhizobia: Multiple facets of competition before, during, and after symbiosis with legumes. Current Opinion in Microbiology, 72, 102281. 10.1016/j.mib.2023.102281

Callahan, B. J., McMurdie, P. J., Rosen, M. J., Han, A. W., Johnson, A. J. A., & Holmes, S. P. (2016). DADA2: High-resolution sample inference from Illumina amplicon data. Nature Methods, 13(7), 581–583. 10.1038/nmeth.3869

Carelli, M., Gnocchi, S., Fancelli, S., Mengoni, A., Paffetti, D., Scotti, C., & Bazzicalupo, M. (2000). Genetic Diversity and Dynamics of *Sinorhizobium meliloti* Populations Nodulating Different Alfalfa Cultivars in Italian Soils. Applied and Environmental Microbiology, 66(11), 4785–4789. 10.1128/AEM.66.11.4785-4789.2000

Chebotar, V. K., Asis, C. A., & Akao, S. (2001). Production of growth-promoting substances and high colonization ability of rhizobacteria enhance the nitrogen fixation of soybean when coinoculated with *Bradyrhizobium japonicum*. Biology and Fertility of Soils, 34(6), 427–432. 10.1007/s00374-001-0426-4

Checcucci, A., DiCenzo, G. C., Bazzicalupo, M., & Mengoni, A. (2017). Trade, Diplomacy, and Warfare: The Quest for Elite Rhizobia Inoculant Strains. Frontiers in Microbiology, 8. 10.3389/fmicb.2017.02207

Cloutier, M. L., Murrell, E., Barbercheck, M., Kaye, J., Finney, D., García-González, I., & Bruns, M. A. (2020). Fungal community shifts in soils with varied cover crop treatments and edaphic properties. Scientific Reports, 10(1), 6198. 10.1038/s41598-020-63173-7

Cole, J. R., Wang, Q., Fish, J. A., Chai, B., McGarrell, D. M., Sun, Y., Brown, C. T., Porras-Alfaro, A., Kuske, C. R., & Tiedje, J. M. (2014). Ribosomal Database Project: Data and tools for high throughput rRNA analysis. Nucleic Acids Research, 42(Database issue), D633–D642. 10.1093/nar/gkt1244

Delestre, C., Laugraud, A., Ridgway, H., Ronson, C., O’Callaghan, M., Barrett, B., Ballard, R., Griffiths, A., Young, S., Blond, C., Gerard, E., & Wakelin, S. (2015). Genome sequence of the clover symbiont *Rhizobium leguminosarum* bv. *trifolii* strain CC275e. Standards in Genomic Sciences, 10(1), 121. 10.1186/s40793-015-0110-1

Denison, R. F., & Kiers, E. T. (2004). Lifestyle alternatives for rhizobia: Mutualism, parasitism, and forgoing symbiosis. FEMS Microbiology Letters, 237(2), 187–193. 10.1111/j.1574-6968.2004.tb09695.x

Denison, R. F., & Kiers, E. T. (2011). Life Histories of Symbiotic Rhizobia and Mycorrhizal Fungi. Current Biology, 21(18), R775–R785. 10.1016/j.cub.2011.06.018

Eddy, S. R. (2011). Accelerated Profile HMM Searches. PLOS Computational Biology, 7(10), e1002195. 10.1371/journal.pcbi.1002195

Efstathiadou, E., Ntatsi, G., Savvas, D., & Tampakaki, A. P. (2021). Genetic characterization at the species and symbiovar level of indigenous rhizobial isolates nodulating *Phaseolus vulgaris* in Greece. Scientific Reports, 11(1), 8674. 10.1038/s41598-021-88051-8

Fernandes, A. D., Macklaim, J. M., Linn, T. G., Reid, G., & Gloor, G. B. (2013). ANOVA-Like Differential Expression (ALDEx) Analysis for Mixed Population RNA-Seq. PLOS ONE, 8(7), e67019. 10.1371/journal.pone.0067019

Finney, D. M., Murrell, E. G., White, C. M., Baraibar, B., Barbercheck, M. E., Bradley, B. A., Cornelisse, S., Hunter, M. C., Kaye, J. P., Mortensen, D. A., Mullen, C. A., & Schipanski, M. E. (2017). Ecosystem services and disservices are bundled in simple and diverse cover cropping systems. Agricultural & Environmental Letters, 2(1), 170033. 10.2134/ael2017.09.0033

Gaby, J. C., & Buckley, D. H. (2012). A Comprehensive Evaluation of PCR Primers to Amplify the *nifH* Gene of Nitrogenase. PLOS ONE, 7(7), e42149. 10.1371/journal.pone.0042149

Galloway, J. N., Townsend, A. R., Erisman, J. W., Bekunda, M., Cai, Z., Freney, J. R., Martinelli, L. A., Seitzinger, S. P., & Sutton, M. A. (2008). Transformation of the Nitrogen Cycle: Recent Trends, Questions, and Potential Solutions. Science, 320(5878), 889–892. 10.1126/science.1136674

Gentleman R, Carey V, Huber W, Hahne F. 2023. _genefilter: genefilter: methods for filtering genes from high-throughput experiments_. doi:10.18129/B9.bioc.genefilter. R package version 1.84.0, https://bioconductor.org/packages/genefilter/.

Ghosh, D., Shi, Y., Zimmermann, I. M., Stürzebecher, T., Holzhauser, K., von Bergen, M., Kaster, A.-K., Spielvogel, S., Dippold, M. A., Müller, J. A., & Jehmlich, N. (2024). Cover crop monocultures and mixtures enhance bacterial abundance and functionality in the maize root zone. ISME Communications, 4(1), ycae132. 10.1093/ismeco/ycae132

Gloor, G. B., Macklaim, J. M., Pawlowsky-Glahn, V., & Egozcue, J. J. (2017). Microbiome Datasets Are Compositional: And This Is Not Optional. Frontiers in Microbiology, 8. 10.3389/fmicb.2017.02224

Gopalakrishnan, S., Sathya, A., Vijayabharathi, R., Varshney, R. K., Gowda, C. L. L., & Krishnamurthy, L. (2015). Plant growth promoting rhizobia: Challenges and opportunities. 3 Biotech, 5(4), 355–377. 10.1007/s13205-014-0241-x

Herridge, D. F., Peoples, M. B., & Boddey, R. M. (2008). Global inputs of biological nitrogen fixation in agricultural systems. Plant and Soil, 311(1), 1–18. 10.1007/s11104-008-9668-3

Hu, L., Busby, R. R., Gebhart, D. L., & Yannarell, A. C. (2014). Invasive *Lespedeza cuneata* and native *Lespedeza virginica* experience asymmetrical benefits from rhizobial symbionts. Plant and Soil, 384(1), 315–325. 10.1007/s11104-014-2213-7

Huang, R., Snedden, W. A., & diCenzo, G. C. (2022). Reference nodule transcriptomes for *Melilotus officinalis* and *Medicago sativa* cv. Algonquin. Plant Direct, 6(6), e408. 10.1002/pld3.408

Huang, X., Zhou, W., Zhang, Y., Yang, Q., Yang, B., Liang, T., Ling, J., & Dong, J. (2024). Keystone PGPR ecological effect: An inoculation case study of diazotrophic *Novosphingobium* sp. N034 on mangrove plant *Kandelia obovate*. Applied Soil Ecology, 202, 105567. 10.1016/j.apsoil.2024.105567

Kaminsky, L. M., Trexler, R. V., Malik, R. J., Hockett, K. L., & Bell, T. H. (2019). The inherent conflicts in developing soil microbial inoculants. Trends in Biotechnology, 37(2), 140–151. 10.1016/j.tibtech.2018.11.011

Kato, N. N., Giongo, A., Zamberlan, P. M., Vahabi, K., Barman, M., Rolli, E., Grosch, R., Lopes, N. P. P., van Dam, N. M. M., & Bueno, P. C. P. (2026). Assessing the chemical composition and ecological relevance of root exudates in legume species. Frontiers in Ecology and Evolution, 14. 10.3389/fevo.2026.1770196

Kaye, J., Finney, D., White, C., Bradley, B., Schipanski, M., Alonso-Ayuso, M., Hunter, M., Burgess, M., & Mejia, C. (2019). Managing nitrogen through cover crop species selection in the U.S. mid-Atlantic. PLOS ONE, 14(4), e0215448. 10.1371/journal.pone.0215448

Kosmopoulos, J. C., Batstone-Doyle, R. T., & Heath, K. D. (2024). Co-inoculation with novel nodule-inhabiting bacteria reduces the benefits of legume–rhizobium symbiosis. Canadian Journal of Microbiology, 70(7), 275–288. 10.1139/cjm-2023-0209

Kuznetsova, A., Brockhoff, P. B., & Christensen, R. H. B. (2017). lmerTest Package: Tests in Linear Mixed Effects Models. Journal of Statistical Software, 82, 1–26. 10.18637/jss.v082.i13

Lenth, R. (2024). emmeans: Estimated Marginal Means, aka Least-Squares Means. R package version 1.10.4, https://rvlenth.github.io/emmeans/, https://rvlenth.github.io/emmeans/

Liebman, A. M., Grossman, J., Brown, M., Wells, M. S., Reberg-Horton, S. C., & Shi, W. (2018). Legume Cover Crops and Tillage Impact Nitrogen Dynamics in Organic Corn Production. Agronomy Journal, 110(3), 1046–1057. 10.2134/agronj2017.08.0474

Martin, M. (2011). Cutadapt removes adapter sequences from high-throughput sequencing reads. EMBnet.journal 17:10.

Martínez-Hidalgo, P., & Hirsch, A. M. (2017). The Nodule Microbiome: N2-Fixing Rhizobia Do Not Live Alone. Phytobiomes Journal, 1(2), 70–82. 10.1094/PBIOMES-12-16-0019-RVW

Maxwell, C. A., Hartwig, U. A., Joseph, C. M., & Phillips, D. A. (1989). A Chalcone and Two Related Flavonoids Released from Alfalfa Roots Induce nod Genes of *Rhizobium meliloti*. Plant Physiology, 91(3), 842–847. 10.1104/pp.91.3.842

McMurdie, P. J., & Holmes, S. (2013). phyloseq: An R Package for Reproducible Interactive Analysis and Graphics of Microbiome Census Data. PLOS ONE, 8(4), e61217. 10.1371/journal.pone.0061217

Mistry, J., Chuguransky, S., Williams, L., Qureshi, M., Salazar, G. A., Sonnhammer, E. L. L., Tosatto, S. C. E., Paladin, L., Raj, S., Richardson, L. J., Finn, R. D., & Bateman, A. (2021). Pfam: The protein families database in 2021. Nucleic Acids Research, 49(D1), D412–D419. 10.1093/nar/gkaa913

Morando, M., Magasin, J. D., Cheung, S., Mills, M. M., Zehr, J. P., & Turk-Kubo, K. A. (2025). Global biogeography of N_2_-fixing microbes: *nifH* amplicon database and analytics workflow. Earth System Science Data, 17(2), 393–422. 10.5194/essd-17-393-2025

Moynihan, M.A., & Furbo Reeder, C. (2023). moyn413/nifHdada2: v2.0.5 (v2.0.5) [Data set]. Zenodo. 10.5281/zenodo.7996213

Mutch, L. A., & Young, J. P. W. (2004). Diversity and specificity of *Rhizobium leguminosarum* biovar *viciae* on wild and cultivated legumes. Molecular Ecology, 13(8), 2435–2444. 10.1111/j.1365-294X.2004.02259.x

Oksanen, J. F., Blanchet, G., Friendly, M., Kindt, R., Legendre, P., McGlinn, D., Minchin, P. R., O’Hara, R. B., Simpson, G. L., Solymos, P., et al. (2019). Vegan: Community Ecology Package. https://cran.r-project.org/package=vegan

Pahua, V. J., Stokes, P. J. N., Hollowell, A. C., Regus, J. U., Gano-Cohen, K. A., Wendlandt, C. E., Quides, K. W., Lyu, J. Y., & Sachs, J. L. (2018). Fitness variation among host species and the paradox of ineffective rhizobia. Journal of Evolutionary Biology, 31(4), 599–610. 10.1111/jeb.13249

Parada, A. E., Needham, D. M., & Fuhrman, J. A. (2016). Every base matters: Assessing small subunit rRNA primers for marine microbiomes with mock communities, time series and global field samples. Environmental Microbiology, 18(5), 1403–1414. 10.1111/1462-2920.13023

Parr, M., Grossman, J. M., Reberg-Horton, S. C., Brinton, C., & Crozier, C. (2011). Nitrogen Delivery from Legume Cover Crops in No-Till Organic Corn Production. Agronomy Journal, 103(6), 1578–1590. 10.2134/agronj2011.0007

Peters, N. K., Frost, J. W., & Long, S. R. (1986). A plant flavone, luteolin, induces expression of *Rhizobium meliloti* nodulation genes. Science, 233(4767), 977–980. 10.1126/science.3738520

Poole, P., Ramachandran, V., & Terpolilli, J. (2018). Rhizobia: From saprophytes to endosymbionts. Nature Reviews Microbiology, 16(5), 291–303. 10.1038/nrmicro.2017.171

Porter, S. S., Dupin, S. E., Denison, R. F., Kiers, E. T., & Sachs, J. L. (2024). Host-imposed control mechanisms in legume–rhizobia symbiosis. Nature Microbiology, 9(8), 1929–1939. 10.1038/s41564-024-01762-2

Rangjaroen, C., Rerkasem, B., Teaumroong, N., Noisangiam, R., & Lumyong, S. (2015). Promoting plant growth in a commercial rice cultivar by endophytic diazotrophic bacteria isolated from rice landraces. Annals of Microbiology, 65(1), 253–266. 10.1007/s13213-014-0857-4

Rho, M., Tang, H., & Ye, Y. (2010). FragGeneScan: Predicting genes in short and error-prone reads. Nucleic Acids Research, 38(20), e191. 10.1093/nar/gkq747

Rocha, R., Lopes, T., Fidalgo, C., Alves, A., Cardoso, P., & Figueira, E. (2023). Bacteria Associated with the Roots of Common Bean (*Phaseolus vulgaris* L.) at Different Development Stages: Diversity and Plant Growth Promotion. Microorganisms, 11(1), 57. 10.3390/microorganisms11010057

Rome, S., Fernandez, M. P., Brunel, B., Normand, P., & Cleyet-Marel, J.-C. (1996). *Sinorhizobium medicae* sp. nov., Isolated from Annual *Medicago* spp. International Journal of Systematic and Evolutionary Microbiology, 46(4), 972–980. 10.1099/00207713-46-4-972

Seitz, V. A., Chaparro, J. M., Schipanski, M. E., Wrighton, K. C., & Prenni, J. E. (2023). Cover crop cultivar, species, and functional diversity is reflected in variable root exudation composition. Journal of Agricultural and Food Chemistry, 71(30), 11373–11385. 10.1021/acs.jafc.3c02912

Seitz, V. A., McGivern, B. B., Borton, M. A., Chaparro, J. M., Schipanski, M. E., Prenni, J. E., & Wrighton, K. C. (2024). Cover crop root exudates impact soil microbiome functional trajectories in agricultural soils. Microbiome, 12(1), 183. 10.1186/s40168-024-01886-x

Shamseldin, A., & Velázquez, E. (2020). The promiscuity of *Phaseolus vulgaris* L. (common bean) for nodulation with rhizobia: A review. World Journal of Microbiology and Biotechnology, 36(5), 63. 10.1007/s11274-020-02839-w

Silva, C., Kan, F. L., & Martínez-Romero, E. (2007). Population genetic structure of *Sinorhizobium meliloti* and *S. medicae* isolated from nodules of *Medicago* spp. In Mexico. FEMS Microbiology Ecology, 60(3), 477–489. 10.1111/j.1574-6941.2007.00301.x

Simbine, M. G., Mohammed, M., Jaiswal, S. K., & Dakora, F. D. (2021). Functional and genetic diversity of native rhizobial isolates nodulating cowpea (*Vigna unguiculata* L. Walp.) in Mozambican soils. Scientific Reports, 11, 12747. 10.1038/s41598-021-91889-7

Smith, G. R., Knight, W. E., Peterson, H. L., & Hagedorn, C. (1982). The Effect of *Rhizobium trifolii* Strains and Crimson Clover Genotypes on N2 Fixation. Crop Science, 22(5), cropsci1982.0011183X002200050017x. 10.2135/cropsci1982.0011183X002200050017x

Steenkamp, E. T., Stępkowski, T., Przymusiak, A., Botha, W. J., & Law, I. J. (2008). Cowpea and peanut in southern Africa are nodulated by diverse *Bradyrhizobium* strains harboring nodulation genes that belong to the large pantropical clade common in Africa. Molecular Phylogenetics and Evolution, 48(3), 1131–1144. 10.1016/j.ympev.2008.04.032

Stewart, J. D., Klein, M., Jaupitre, S., Oyarte-Galvez, L., Dong, L., Bouwmeester, H. J., Kiers, E. T., Kokkoris, V., & Weedon, J. T. (2026). Root traits correlate with crop rhizosphere microbiome diversity independent of legume relatedness. ISME Communications, 6(1), ycag087. 10.1093/ismeco/ycag087

Stopnisek, N., & Shade, A. (2021). Persistent microbiome members in the common bean rhizosphere: An integrated analysis of space, time, and plant genotype. The ISME Journal, 15(9), 2708–2722. 10.1038/s41396-021-00955-5

Tao, K., Jensen, I. T., Zhang, S., Villa-Rodríguez, E., Blahovska, Z., Salomonsen, C. L., Martyn, A., Björgvinsdóttir, Þ. N., Kelly, S., Janss, L., Glasius, M., Waagepetersen, R., & Radutoiu, S. (2024). Nitrogen and Nod factor signaling determine *Lotus japonicus* root exudate composition and bacterial assembly. Nature Communications, 15(1), 3436. 10.1038/s41467-024-47752-0

Thilakarathna, M. S., & Raizada, M. N. (2017). A meta-analysis of the effectiveness of diverse rhizobia inoculants on soybean traits under field conditions. Soil Biology and Biochemistry, 105, 177–196. 10.1016/j.soilbio.2016.11.022

Thrall, P. H., Laine, A.-L., Broadhurst, L. M., Bagnall, D. J., & Brockwell, J. (2011). Symbiotic Effectiveness of Rhizobial Mutualists Varies in Interactions with Native Australian Legume Genera. PLOS ONE, 6(8), e23545. 10.1371/journal.pone.0023545

Wagner, M. R., Busby, P. E., & Balint-Kurti, P. (2020). Analysis of leaf microbiome composition of near-isogenic maize lines differing in broad-spectrum disease resistance. New Phytologist, 225(5), 2152–2165. 10.1111/nph.16284

Walker, L., Lagunas, B., & Gifford, M. L. (2020). Determinants of host range specificity in legume-rhizobia symbiosis. Frontiers in Microbiology, 11, 585749. 10.3389/fmicb.2020.585749

Wickham, H., Averick, M., Bryan, J., Chang, W., McGowan, L. D., François, R., Grolemund, G., Hayes, A., Henry, L., Hester, J., Kuhn, M., Pedersen, T. L., Miller, E., Bache, S. M., Müller, K., Ooms, J., Robinson, D., Seidel, D. P., Spinu, V., … Yutani, H. (2019). Welcome to the Tidyverse. Journal of Open Source Software, 4(43), 1686. 10.21105/joss.01686

Yan, J., Han, X. Z., Ji, Z. J., Li, Y., Wang, E. T., Xie, Z. H., & Chen, W. F. (2014). Abundance and Diversity of Soybean-Nodulating Rhizobia in Black Soil Are Impacted by Land Use and Crop Management. Applied and Environmental Microbiology, 80(17), 5394–5402. 10.1128/AEM.01135-14

Yao, Z. Y., Kan, F. L., Wang, E. T., Wei, G. H., & Chen, W. X. (2002). Characterization of rhizobia that nodulate legume species of the genus *Lespedeza* and description of *Bradyrhizobium yuanmingense* sp. nov. International Journal of Systematic and Evolutionary Microbiology, 52(6), 2219–2230. 10.1099/00207713-52-6-2219

Yu, Y.-H., Crosbie, D. B., & Arancibia, M. M. (2025). *Pseudomonas* in the spotlight: Emerging roles in the nodule microbiome. Trends in Plant Science, 30(5), 461–470. 10.1016/j.tplants.2024.12.002

