## Supplemental materials for "Cover crop microbiomes affect legume cash crop growth but not consistently through enriching nitrogen-fixing rhizobia"

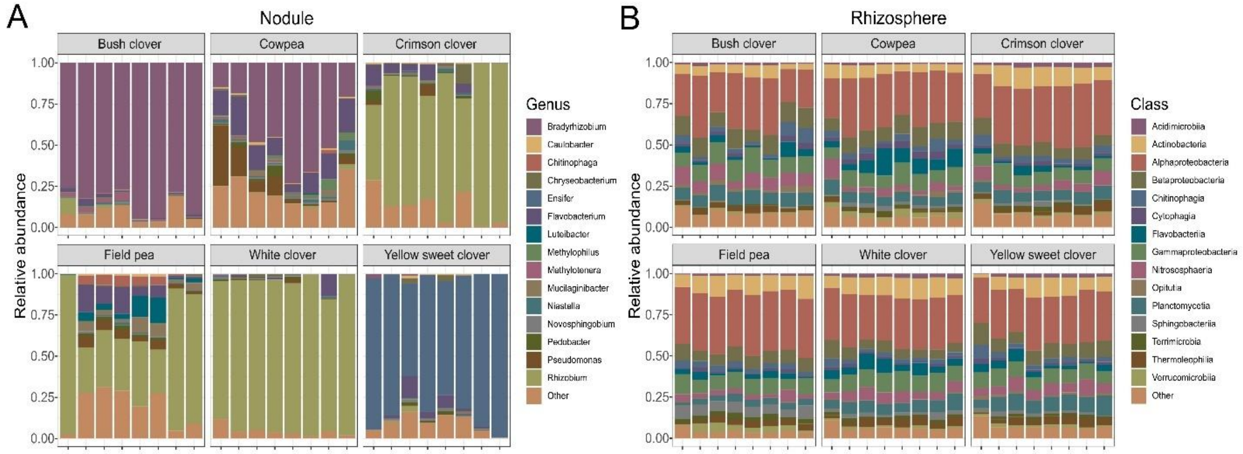


**Supplemental Fig. 1.** Bacterial community composition of cover crop (A) nodules and (B) rhizospheres across replicates assessed by 16S rRNA amplicon sequencing. The top 15 genera and top 15 classes are shown for nodules and rhizospheres, respectively, with remaining taxa collapsed into 'Other'."

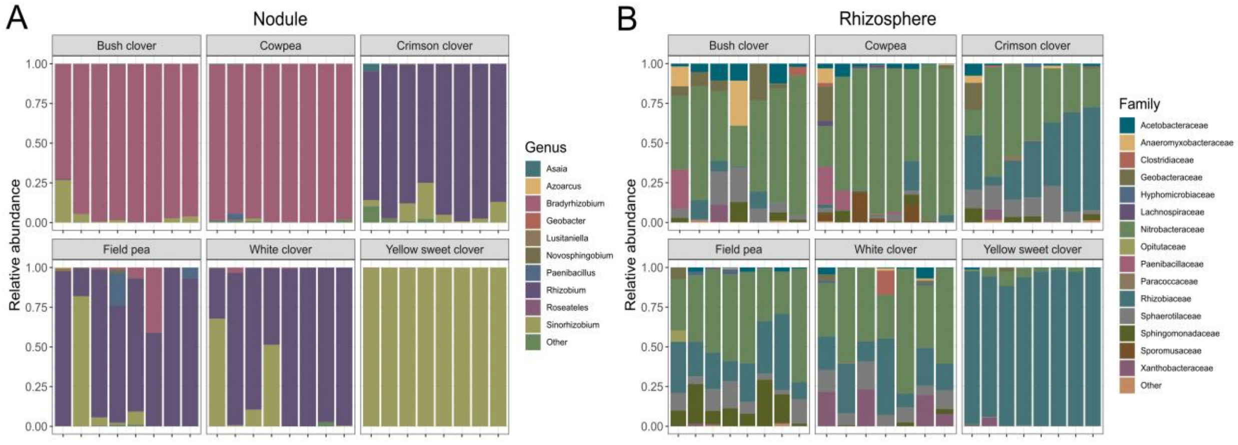


**Supplemental Fig. 2.** Diazotroph community composition of cover crop (A) nodules and (B) rhizospheres across replicates assessed by *nifH* amplicon sequencing. The top 10 genera and top 15 families are shown for nodules and rhizospheres, respectively, with remaining taxa collapsed into 'Other'.

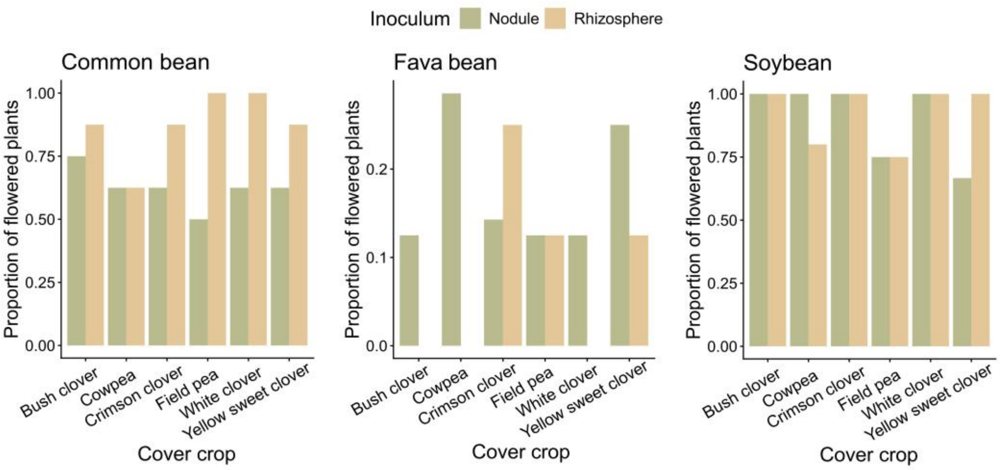


**Supplemental Fig. 3.** The effect of cover crop inocula on the proportion of flowered cash crops after approximately 12 weeks of growth. Alfalfa is excluded since no plants flowered.

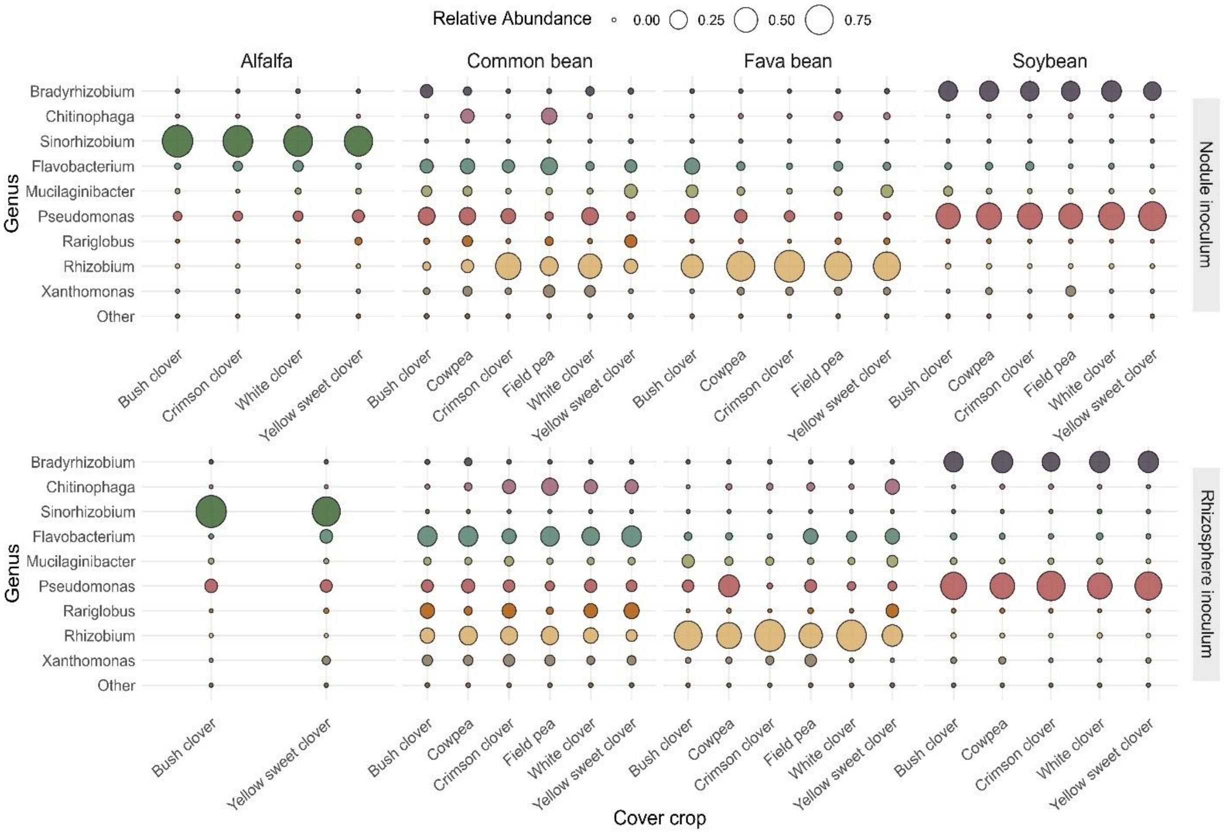

**Supplemental Fig. 4.** Bacterial community composition of cash crop nodules receiving cover crop nodule or rhizosphere inoculum, assessed by 16S rRNA amplicon sequencing. The nine most abundant genera (pooled across samples) are shown. Remaining taxa are combined into 'Other'.

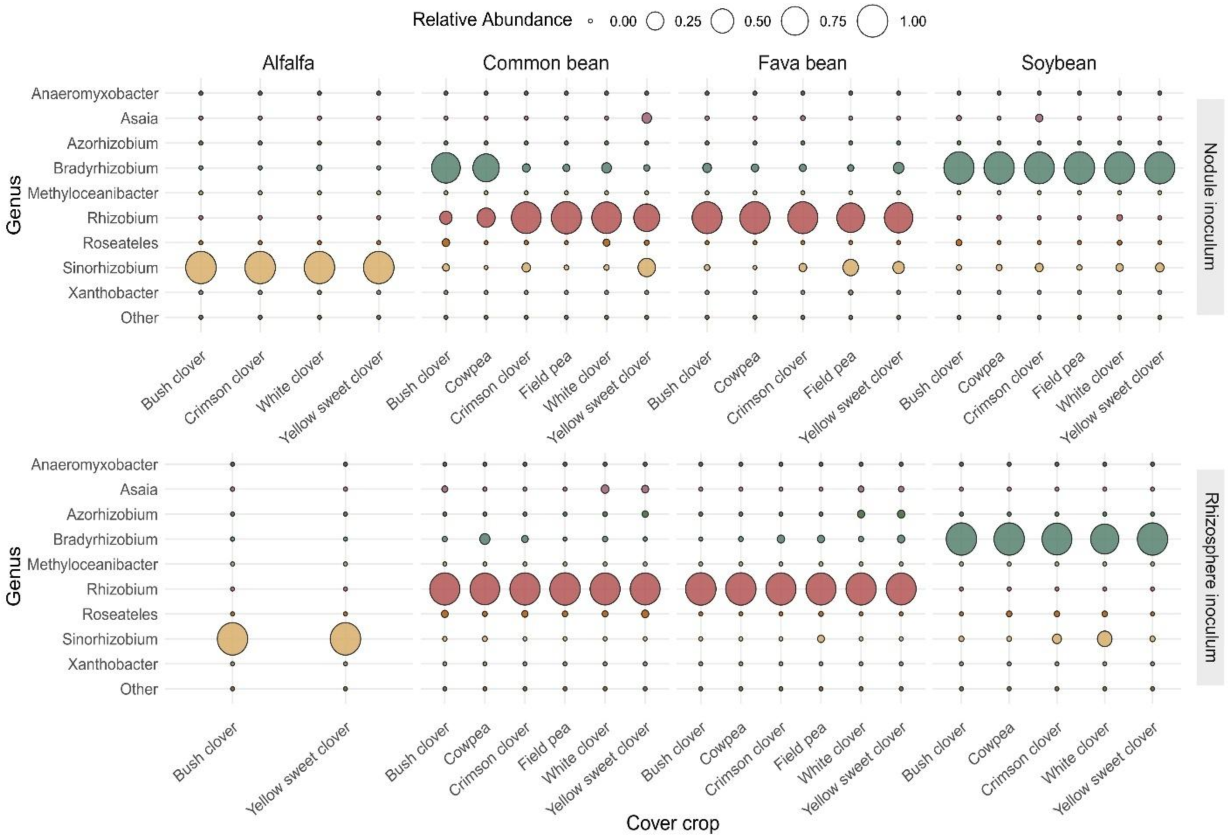


**Supplemental Fig. 5.** Diazotroph community composition of cash crop nodules that received cover crop nodule or rhizosphere inoculum assessed by *nifH* amplicon sequencing. The nine most abundant genera (pooled across samples) are shown with remaining taxa collapsed into 'Other'.

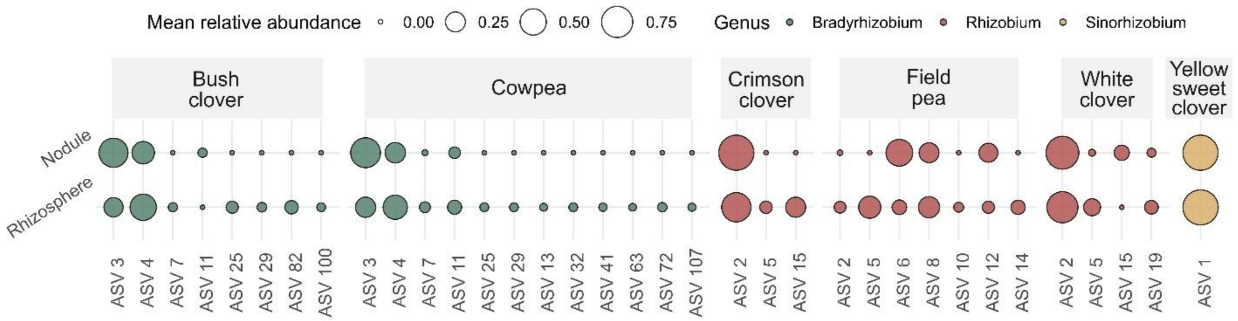


**Supplemental Fig. 6.** ASV-level composition of the *nifH* rhizobial symbiont genus associated with each cover crop (*Bradyrhizobium* for bush clover, cowpea; *Rhizobium* for crimson clover, field pea, white clover; *Sinorhizobium* for yellow sweet clover), shown separately for nodule and rhizosphere compartment. For each cover crop, the smallest set of ASVs collectively accounting for over 97% of mean relative abundance across replicates is shown. Bubble size reflects mean relative abundance within each compartment; color indicates genus.

**Supplemental Table 1.** Shannon and species (ASV) richness diversity indices for the rhizosphere and nodule cover crop 16S communities. P-values were adjusted using the Benjamini-Hochberg method to correct for multiple comparisons.

| **Response** | **Term** | **Numerator Degrees of Freedom** | **Denominator Degrees of Freedom** | **F-value** | **P-adjusted** |
|---|---|---|---|---|---|
| Shannon diversity | Compartment | 1 | 78.37 | 947.62 | **2.37E-45** |
| Shannon diversity | Cover crop | 5 | 75.78 | 12.20 | **3.31E-08** |
| Shannon diversity | Compartment x cover crop | 5 | 76.72 | 13.36 | **6.84E-09** |
| Shannon diversity | Sequencing depth | 1 | 79.89 | 0.05 | 9.31E-01 |
| Species richness | Compartment | 1 | 78.69 | 956.43 | **2.37E-45** |
| Species richness | Cover crop | 5 | 75.61 | 9.99 | **3.45E-07** |
| | | 5 | 76.72 | 6.42 | **4.91E-05** |

| | | | | | |
|---|---|---|---|---|---|
| Species richness | Compartment x cover crop | | | | |
| Species richness | Sequencing depth | 1 | 76.79 | 16.01 | **4.33E-04** |

**Supplemental Table 2.** Shannon and species (ASV) richness diversity indices for the rhizosphere and nodule cover crop *nifH* communities. P-values were adjusted using the Benjamini-Hochberg method to correct for multiple comparisons.

| **Response** | **Term** | **Numerator Degrees of Freedom** | **Denominator Degrees of Freedom** | **F-value** | **P-adjusted** |
|---|---|---|---|---|---|
| Shannon diversity | Compartment | 1 | 78.89 | 15.56 | **2.59E-04** |
| Shannon diversity | Cover crop | 5 | 74.47 | 3.82 | **1.17E-02** |
| Shannon diversity | Compartment x cover crop | 5 | 74.01 | 5.02 | **1.53E-03** |
| Shannon diversity | Sequencing depth | 1 | 79.65 | 0.74 | 4.04E-01 |
| Species richness | Compartment | 1 | 78.05 | 35.03 | **2.46E-07** |
| Species richness | Cover crop | 5 | 74.18 | 2.31 | **5.27E-02** |
| Species richness | Compartment x cover crop | 5 | 73.8 | 1.26 | 2.91E-01 |
| Species richness | Sequencing depth | 1 | 78.92 | 2.23 | 4.04E-01 |

**Supplemental Table 3.** Relative abundance of the known rhizobial symbiont genus associated with each cover crop species in nodule communities determined by 16S rRNA and *nifH* amplicon sequencing. The number of ASVs assigned to the rhizobial symbiont genus and their mean relative abundance (± SD) across replicates are reported for each cover crop.

| Cover crop | Genus | Amplicon | Number of ASVs | Mean relative abundance | Mean relative abundance (SD) |
|---|---|---|---|---|---|
| Bush clover | *Bradyrhizobium* | 16S | 2 | 0.822 | 0.094 |
| Cowpea | *Bradyrhizobium* | 16S | 2 | 0.405 | 0.213 |
| Crimson clover | *Rhizobium* | 16S | 4 | 0.740 | 0.199 |
| Field pea | *Rhizobium* | 16S | 3 | 0.523 | 0.295 |
| White clover | *Rhizobium* | 16S | 3 | 0.904 | 0.062 |
| Yellow sweet clover | *Sinorhizobium* | 16S | 1 | 0.815 | 0.140 |
| Bush clover | *Bradyrhizobium* | *nifH* | 4 | 0.946 | 0.091 |
| Cowpea | *Bradyrhizobium* | *nifH* | 6 | 0.983 | 0.020 |
| Crimson clover | *Rhizobium* | *nifH* | 5 | 0.897 | 0.087 |
| Field pea | *Rhizobium* | *nifH* | 7 | 0.771 | 0.278 |
| White clover | *Rhizobium* | *nifH* | 7 | 0.826 | 0.270 |
| Yellow sweet clover | *Sinorhizobium* | *nifH* | 13 | 0.999 | 0.002 |

**Supplemental Table 4.** Analysis of variance (ANOVA) was performed on square-root transformed nodule counts using linear models with Type III sums of squares. Significance of model terms was assessed using F-tests.

| | *Alfalfa* | | *Common bean* | | *Fava bean* | | *Soybean* | |
|---|---|---|---|---|---|---|---|---|
| Compartment | $F_{1,59}$= 1.59 | P = 0.212 | $F_{1,84}$= 0.04 | P = 0.849 | $F_{1,72}$= 0.95 | P = 0.332 | $F_{1,43}$= 0.46 | P = 0.458 |
| Cover crop | $F_{5,59}$= 12.1 | **P < 0.001** | $F_{5,84}$= 2.73 | **P = 0.025** | $F_{5,72}$= 5.9 | **P < 0.001** | $F_{5,43}$= 3.58 | **P = 0.009** |
| Compartment x cover crop | $F_{5,59}$= 1.77 | P = 0.133 | $F_{5,84}$= 2.89 | **P = 0.019** | $F_{5,72}$= 1.96 | P = 0.095 | $F_{5,43}$= 0.52 | P = 0.756 |

**Supplemental Table 5.** Analysis of variance (ANOVA) was performed on total plant biomass using linear models with Type III sums of squares. Alfalfa total biomass was log-transformed prior to analysis to meet model assumptions. Significance of model terms was assessed using F-tests.

| | *Alfalfa* | | *Common bean* | | *Fava bean* | | *Soybean* | |
|---|---|---|---|---|---|---|---|---|
| Compartment | $F_{1,63}$= 0.15 | P = 0.698 | $F_{1,77}$= 19.3 | **P < 0.001** | $F_{1,72}$= 1.37 | P = 0.245 | $F_{1,40}$= 0.54 | P = 0.467 |

| | | | | | | | | |
|---|---|---|---|---|---|---|---|---|
| Cover crop | $F_{5,63}$= 8.11 | **P < 0.001** | $F_{5,77}$= 1.72 | P = 0.139 | $F_{5,72}$= 2.39 | **P = 0.046** | $F_{5,40}$= 0.31 | P = 0.905 |
| Compartment x cover crop | $F_{5,63}$= 0.27 | P = 0.931 | $F_{5,77}$= 1.29 | P = 0.275 | $F_{5,72}$= 0.53 | P = 0.782 | $F_{5,40}$= 0.75 | P = 0.594 |

**Supplemental Table 6.** Logistic regression analysis of proportion of flowered plants for each cash crop. Alfalfa is excluded since no plants flowered. Likelihood ratio tests were used for significance testing.

| | *Common bean* | | *Fava bean* | | *Soybean* | |
|---|---|---|---|---|---|---|
| Compartment | $\chi^2_1$ = 8.29 | **P = 0.004** | $\chi^2_1$ = 1.56 | P = 0.212 | $\chi^2_1$ = 0.07 | P = 0.791 |
| Cover crop | $\chi^2_5$ = 2.12 | P = 0.833 | $\chi^2_5$ = 2.57 | P = 0.765 | $\chi^2_5$ = 7.09 | P = 0.214 |
| Compartment x cover crop | $\chi^2_5$ = 6.46 | P = 0.264 | $\chi^2_5$ = 4.86 | P = 0.433 | $\chi^2_5$ = 3.69 | P = 0.595 |

**Supplemental Table 7.** Analysis of variance (ANOVA) for linear mixed-effects models testing the effect of inoculum cell count on total biomass and nodule number for each cash crop. F-tests were used for significance testing.

| | *Alfalfa* | | *Common bean* | | *Fava bean* | | *Soybean* | |
|---|---|---|---|---|---|---|---|---|
| Nodule number | $F_{1,73}$= 0.01 | P = 0.910 | $F_{1,94}$= 1.70 | P = 0.195 | $F_{1,89}$= 0.46 | P = 0.501 | $F_{1,50}$= 0.10 | P = 0.292 |
| Total biomass | $F_{1,50}$= 0.10 | P = 0.754 | $F_{1,94}$= 0.01 | P = 0.913 | $F_{1,86}$= 3.64 | P = 0.060 | $F_{1,51}$= 1.13 | P = 0.753 |

**Supplemental Table 8.** Relative abundance of rhizobial genera associated with cash crop species in nodule communities determined by 16S rRNA amplicon sequencing. The number of ASVs assigned to the rhizobial genus and their mean relative abundance (± SD) across cover crops for nodule or rhizosphere inoculum are reported for each cash crop.

| Cash crop | Inoculum | Genus | Number of ASVs | Mean relative abundance | Mean relative abundance (SD) |
|---|---|---|---|---|---|
| Alfalfa | Nodule | *Bradyrhizobium* | 1 | 0.00018 | 0.00021 |
| Alfalfa | Rhizosphere | *Bradyrhizobium* | 1 | 0.00013 | 0.00005 |
| Alfalfa | Nodule | *Rhizobium* | 2 | 0.00036 | 0.00029 |
| Alfalfa | Rhizosphere | *Rhizobium* | 4 | 0.00019 | 0.00005 |
| Alfalfa | Nodule | *Sinorhizobium* | 1 | 0.84726 | 0.05903 |
| Alfalfa | Rhizosphere | *Sinorhizobium* | 1 | 0.79457 | 0.11229 |
| Common bean | Nodule | *Bradyrhizobium* | 2 | 0.02497 | 0.03604 |
| Common bean | Rhizosphere | *Bradyrhizobium* | 1 | 0.00287 | 0.00613 |
| Common bean | Nodule | *Rhizobium* | 6 | 0.25313 | 0.22051 |
| Common bean | Rhizosphere | *Rhizobium* | 4 | 0.16336 | 0.06347 |
| Common bean | Nodule | *Sinorhizobium* | 1 | 0.00013 | 0.00031 |
| Common bean | Rhizosphere | *Sinorhizobium* | *0* | *0* | *0* |
| Fava bean | Nodule | *Bradyrhizobium* | 1 | 0.00054 | 0.00054 |
| Fava bean | Rhizosphere | *Bradyrhizobium* | 1 | 0.00029 | 0.00022 |
| Fava bean | Nodule | *Rhizobium* | 8 | 0.69570 | 0.17159 |
| Fava bean | Rhizosphere | *Rhizobium* | 5 | 0.61326 | 0.21564 |
| Fava bean | Nodule | *Sinorhizobium* | 1 | 0.00004 | 0.00003 |
| Fava bean | Rhizosphere | *Sinorhizobium* | *0* | *0* | *0* |
| Soybean | Nodule | *Bradyrhizobium* | 2 | 0.28694 | 0.02958 |
| Soybean | Rhizosphere | *Bradyrhizobium* | 2 | 0.31156 | 0.04551 |
| Soybean | Nodule | *Rhizobium* | 6 | 0.00127 | 0.00094 |
| Soybean | Rhizosphere | *Rhizobium* | 6 | 0.00099 | 0.00084 |
| Soybean | Nodule | *Sinorhizobium* | 1 | 0.00005 | 0.00003 |
| Soybean | Rhizosphere | *Sinorhizobium* | 1 | 0.00020 | 0.00037 |

**Supplemental Table 9.** Relative abundance of rhizobial genera associated with cash crop species in nodule communities determined by *nifH* amplicon sequencing. The number of ASVs assigned to the rhizobial genus and their mean relative abundance (± SD) across cover crops for nodule or rhizosphere inoculum are reported for each cash crop.

| Cash crop | Inoculum | Genus | Number of ASVs | Mean relative abundance | Mean relative abundance (SD) |
|---|---|---|---|---|---|
| Alfalfa | Nodule | *Bradyrhizobium* | 2 | 0.00056 | 0.00113 |
| Alfalfa | Rhizosphere | *Bradyrhizobium* | 1 | 0.00003 | 0.00004 |

| | | | | | |
|---|---|---|---|---|---|
| Alfalfa | Nodule | *Rhizobium* | *0* | *0* | *0* |
| Alfalfa | Rhizosphere | *Rhizobium* | *0* | *0* | *0* |
| Alfalfa | Nodule | *Sinorhizobium* | 6 | 0.99930 | 0.00138 |
| Alfalfa | Rhizosphere | *Sinorhizobium* | 9 | 0.99991 | 0.00010 |
| Common bean | Nodule | *Bradyrhizobium* | 8 | 0.27326 | 0.39360 |
| Common bean | Rhizosphere | *Bradyrhizobium* | 8 | 0.01150 | 0.02117 |
| Common bean | Nodule | *Rhizobium* | 14 | 0.65492 | 0.37984 |
| Common bean | Rhizosphere | *Rhizobium* | 11 | 0.96856 | 0.01843 |
| Common bean | Nodule | *Sinorhizobium* | 3 | 0.05016 | 0.10105 |
| Common bean | Rhizosphere | *Sinorhizobium* | 1 | 0.00030 | 0.00056 |
| Fava bean | Nodule | *Bradyrhizobium* | 8 | 0.02501 | 0.01971 |
| Fava bean | Rhizosphere | *Bradyrhizobium* | 10 | 0.00830 | 0.00809 |
| Fava bean | Nodule | *Rhizobium* | 8 | 0.88138 | 0.10564 |
| Fava bean | Rhizosphere | *Rhizobium* | 7 | 0.95605 | 0.04273 |
| Fava bean | Nodule | *Sinorhizobium* | 2 | 0.05580 | 0.07633 |
| Fava bean | Rhizosphere | *Sinorhizobium* | 1 | 0.00208 | 0.00509 |
| Soybean | Nodule | *Bradyrhizobium* | 9 | 0.97342 | 0.01409 |
| Soybean | Rhizosphere | *Bradyrhizobium* | 8 | 0.95481 | 0.07072 |
| Soybean | Nodule | *Rhizobium* | 3 | 0.00094 | 0.00199 |
| Soybean | Rhizosphere | *Rhizobium* | *0* | *0* | *0* |
| Soybean | Nodule | *Sinorhizobium* | 1 | 0.01322 | 0.01132 |
| Soybean | Rhizosphere | *Sinorhizobium* | 2 | 0.04088 | 0.06927 |

104
